# Short-term fluctuating and long-term divergent selection on sympatric Monkeyflowers: insights from repeated reciprocal transplants

**DOI:** 10.1101/2024.06.26.600870

**Authors:** Caroline M. Dong, Bolívar Aponte Rolón, Juj K. Sullivan, Diana Tataru, Max Deleon, Rachael Dennis, Spencer Dutton, Fidel J. Machado Perez, Lissette Montano, Kathleen G. Ferris

## Abstract

Sympatric species are often locally adapted to distinct microhabitats, yet temporal variation may cause local maladaptation and weaken species boundaries, particularly during extreme climatic events. To examine the interplay between spatially and temporally varying selection, we studied sympatric Monkeyflowers occupying dramatically different niches, *Mimulus guttatus* and *M. laciniatus.* We conducted three replicated reciprocal transplants and combined them with previous experiments to leverage a dataset of five single-year transplants spanning a decade. We estimated the strength of phenotypic selection on experimental hybrids in each species’ habitat across years of drastically differing snowpack. If ecological isolation maintains species differences, we predicted divergent phenotypic selection between habitats consistent with species’ differences and local adaptation. We found spatial and interannual fluctuations in phenotypic selection, often in unpredicted directions, with temporal variation associated with snowpack. In a combined-year analysis, we detected a significant difference in the strength of long-term selection on flowering time, a key temporally isolating and adaptative trait, between species’ habitats, suggesting that selection may reinforce species boundaries despite short-term fluctuations. Finally, we found parental local adaptation varied among years where *M. laciniatus* was locally adapted during low snowpack years, while an extreme snowfall event contributed to overall local maladaptation of *M. guttatus*.

## INTRODUCTION

Closely related species often occupy distinct habitats within areas of geographic overlap (Coyne and Orr 2004; Rundle and Nosil 2005; Schluter 2009). Species boundaries in sympatry can be maintained by adaptation to differing environments, generating reproductive isolation via selection against immigrants (e.g. habitat isolation) and intermediate hybrids (e.g. extrinsic post-zygotic isolation; Rundle & Nosil, 2005). Identifying the traits contributing to reproductive isolation is difficult because as speciation progresses, multiple forms of reproductive isolation accumulate which hinders the measurement of individual barriers (Coyne and Orr 2004). Neutral processes also cause the accumulation of phenotypic divergence. Thus, experimental manipulation is necessary to determine whether habitat isolation is a key barrier, and which traits are involved in divergent adaptation (Ramsey et al. 2003; Hall and Willis 2006; Lowry et al. 2008). Measuring phenotypic selection on closely related sympatric species in a reciprocal transplant can identify whether divergent adaptation is important for reproductive isolation and speciation (Lowry et al. 2008; Anderson et al. 2015). If so, we expect each species to have higher fitness in its habitat and phenotypic selection in the direction of species’ differences (Anderson et al. 2015*a*), indicating habitat isolation and local adaptation to current conditions.

Despite these predictions of local adaptation, maladaptation to native habitats is often observed in reciprocal transplants (Hereford 2009; Kooyers et al. 2019; Dittmar and Schemske 2023). This could be due to the prevalence of temporal fluctuations in selection (Siepielski et al. 2009) which can influence the strength of reproductive isolation and species boundaries (Tataru et al. 2023, 2024). Additionally, current environmental conditions may have changed from the conditions of original divergence (Coyne and Orr 2004; Rundle and Nosil 2005). If species are no longer adapted to their current niches due to climate change, then we expect to find local maladaptation (Crespi 2000) and selection for the non-local species’ phenotype (Wilczek et al. 2014; Kooyers et al. 2019; Anderson and Wadgymar 2020). In this scenario, spatially varying selection may act contrary to reproductive isolation and in the absence of strong intrinsic barriers, could erode species boundaries by driving each towards the other’s phenotypic optimum.

Variation in selection in longitudinal studies is usually measured on a seasonal or inter-annual timescale, which is a short-term measurement of population evolution. This leads to the question of how short-term fluctuations affect long-term evolutionary change in species and populations (Wadgymar et al. 2017; Kelly 2022; Dittmar and Schemske 2023; Oakley et al. 2023). A synthesis of phenotypic selection studies by Kingsolver & Diamond (2011) found that while temporal variation in selection is common, it does not significantly inhibit long-term directional selection. Therefore, even with fluctuations in the direction of selection, there may be an average directional or stabilizing trend over a longer timescale. For example, Wadgymar et al. (2017) found that interannual fluctuations in the direction of selection on the perennial plant *Boechera stricta* gave rise to long-term patterns consistent with stabilizing selection and local adaptation. In *Arabidopsis thaliana*, Lee et al. (2024) found long-term divergent selection on cold tolerance between populations in Italy and Sweden despite interannual fluctuations. However, due to the arduousness of temporally replicating reciprocal transplant experiments, it is not currently well understood how short versus longer-term patterns of natural selection compare and impact species’ divergence and reproductive isolation.

Another understudied agent of evolutionary change is episodic selection stemming from sudden and extreme environmental events. Long-term studies capturing the evolutionary responses to these dramatic ecological changes are rare yet critical to understand their impact on species boundaries (Coltman et al. 1999; Pemberton et al. 2022). For instance, through their long-term study of the Galapagos finches, Grant and Grant (1993) were able to illustrate how an extreme El Niño year altered selection on beak size and rates of interspecific gene flow. Strong episodes of selection such as this can impact biodiversity by affecting the evolutionary trajectory of species adaptation and reproductive isolation (Grant and Grant 1993). Studies such as this are vital because extreme climatic events are predicted to become more frequent with continuing anthropogenic climate change (Bailey and van de Pol 2016).

The *Mimulus guttatus* species complex is well-suited for studying how natural selection shapes species boundaries and reproductive isolation (Wu et al. 2008). *Mimulus guttatus* and *M. laciniatus* (syn. *Erythranthe guttata* and *E. laciniata*) occur in contrasting microhabitats throughout central and southern Sierra Nevada, CA. *Mimulus guttatus* inhabits moist meadows and seeps while *M. laciniatus* occupies rocky outcrops characterized by shallow soils, intense light, extreme temperatures, and a short growing season (Ferris et al. 2014; Ferris and Willis 2018). *Mimulus laciniatus*, an obligate summer annual, has a smaller stature, lobed leaves, self-fertilization with reduced floral size, and early rapid flowering (DeMarche et al. 2013; Ferris and Willis 2018). In contrast, *M. guttatus* populations are facultatively annual, later-flowering, predominantly outcrossing, round-leaved, and larger in size and flower morphology (Awadalla and Ritland 1997; Ferris et al. 2014). Despite ecological divergence, reproductive isolation is incomplete (Vickery 1964; Ferris et al. 2017; Tataru et al. 2024).

A previous study demonstrated local adaptation to contrasting microhabitats, but it remains unclear how environmental variation alters the strength or direction of selection over time (Ferris and Willis 2018). Early flowering, a drought escape strategy, functions as a key pre-mating barrier in the *Mimulus guttatus* species complex (Hall and Willis 2006; Lowry et al. 2008; Ferris et al. 2017; Mantel and Sweigart 2019). Lobed leaves may improve temperature and water regulation in rocky habitats (Nicotra et al. 2011*a*). Yet how selection on these and other traits, including plant size, mating system, and flower morphology, fluctuates through time is unknown. Both species exhibit phenotypic plasticity with *M. guttatus* exhibiting greater plasticity in plant size and flowering time than *M. laciniatus* (Ferris & Willis 2018). Such plasticity may influence fitness across habitats, but its contribution to adaptation and reproductive isolation remains unclear.

Here, we investigate spatial and temporal variation in natural selection between *M. guttatus* and *M. laciniatus* using five single-year reciprocal transplants under contrasting climatic conditions (Figure 1). We add three years of new results to two previously published experiments (Ferris and Willis 2018, Tataru et al., 2023) and examine transplants during low snowpack years (2013, 2021, 2022) and high snowpack years (2019, 2023; Figure 1a,b). We address the following questions: (1) How do environmental variables affect fitness in each species’ habitat and induce phenotypic plasticity? (2) Does the strength and direction of selection on adaptive and reproductively isolating phenotypes fluctuate? (3) How do short versus long-term patterns of selection compare? (4) Are *M. guttatus* and *M. laciniatus* locally adapted to their respective microhabitats? (5) How does episodic selection affect the trajectory of species divergence? By examining both interannual variation and overarching signatures of selection, we were able to elucidate the effects of a changing climate on the maintenance of biodiversity in this system.

**Figure 1.**
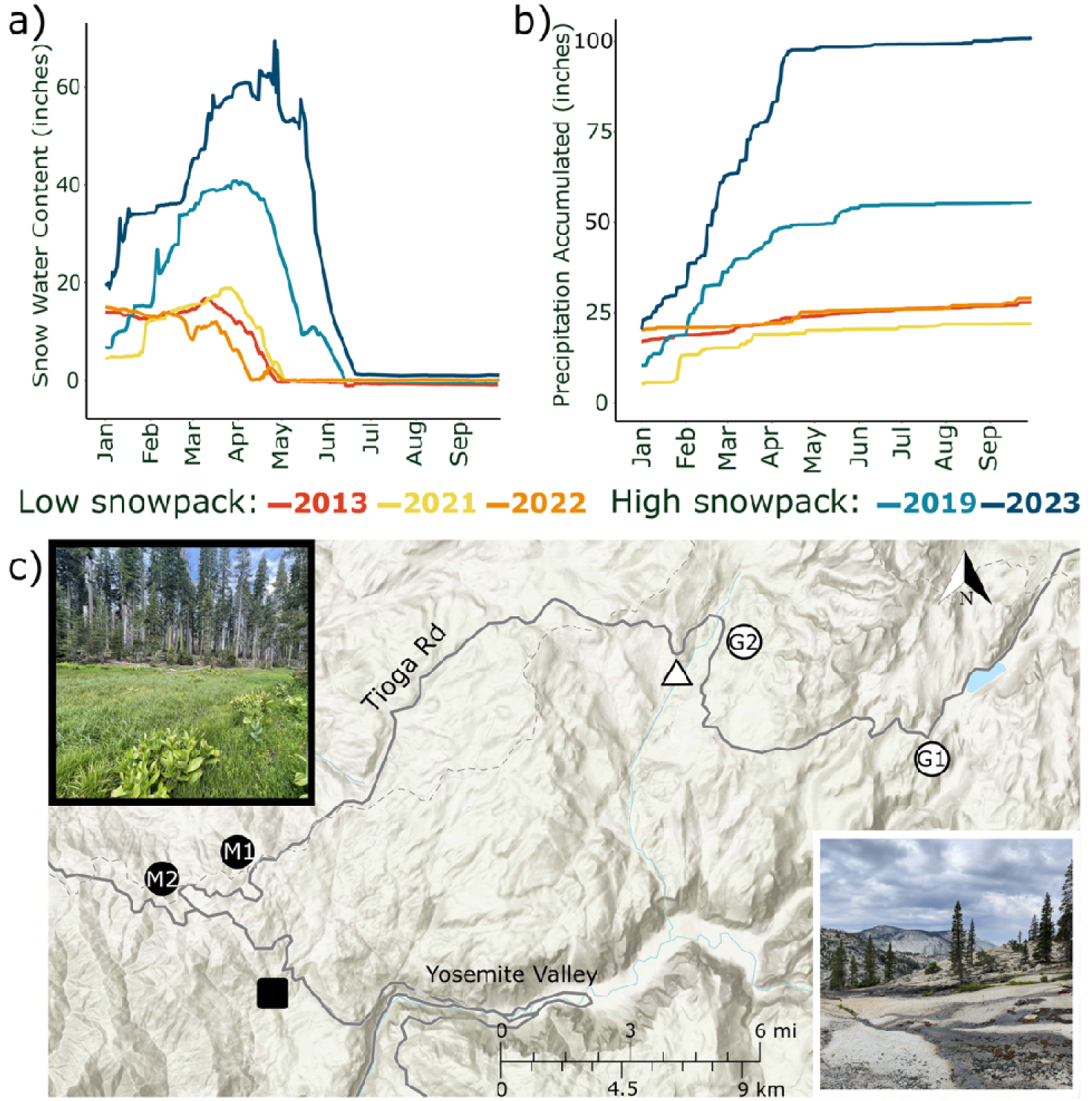
Temporal and spatial scale of the experimental design. Annual changes in a) snow water content and b) accumulated precipitation during experimental years of low snowpack (2013, 2021, 2022) and comparatively higher snowpack (2019 and 2023). Data are averaged from the California Department of Water Resources data stations GIN, WHW, and TUM, located nearby experimental sites along Tioga Road in Yosemite National Park (cdec.water.ca.gov). The c) spatial layout of experimental sites: meadow 1 and 2 (M1 and M2; filled circles) and granite 1 and 2 (G1 and G2; open circles). Nearby parental source populations for experimental crosses are shown: *M*. *guttatus* YVO (filled square) and *M*. *laciniatus* WLF (open triangle). Photos of representative meadow and granite habitats are shown in the insets.

## MATERIALS AND METHODS

### Repeated reciprocal transplant design

Reciprocal transplants used hybrids created from inbred lines of *M. guttatus* (YVO 18; 37.723366, -119.746433; 5,395 ft elevation) and *M. laciniatus* (WLF 47; 37.841533, -119.59385; 7993 ft elevation; Figure 1c). Advanced generation hybrids were used because recombination over multiple generations breaks up parental haplotypes, increasing phenotypic variation and enabling inferences about selection on individual traits (Figure S1; Nagy, 1997; Hall & Willis, 2006; Groh & Coop, 2024). F_1_ hybrids (WLF 47 maternal x YVO 18 paternal) were self-fertilized to generate F_2_ seeds used in 2021. Five hundred F_2_ hybrids were propagated with single-seed descent to produce F_3_ seeds used in 2022 and 300 F_6_ lines used in 2023. Of the initial 500, 179 lines were lost during propagation due to a combination of inbreeding depression and logistical issues. Therefore, the F_6_ hybrid population is a subset of the genetic variation present in the original F_2_ population. In 2021 and 2022, seeds from 200 hybrid maternal families were pooled for transplants. In 2023, parental inbred lines were intercrossed within each parental population to avoid inbreeding depression, which can impact inferences about local adaptation (Willis 1999). The 2013 and 2019 experiments used F_4_ hybrids from the same parental populations (see Ferris & Willis 2018; Tataru et al 2023).

Our transplants were conducted in 2021–2023 in Yosemite National Park, CA, replicating experiments from 2013 and 2019 at four sites (Figure 1c): two granite outcrops (Granite 1 [37.810700, -119.485200; 8,436 ft elevation], Granite 2 [37.843702, -119.573120; 8,117 ft elevation]) and two meadows (Meadow 1 [37.767781, -119.772720; 6,605 ft elevation], Meadow 2 [37.755968, -119.803031; 6,163 ft elevation]). We used the same microhabitats within each site, experimental design, and parental source populations. Sample sizes, maternal families, and hybrid generation differed annually within logistical constraints (Table S1). Seedlings were transplanted in small blocks to capture fine-scale environmental variation and mimic granite habitat patchiness. In 2021 and 2022, each site had 50 randomized blocks (4×6 formation, 1-inch spacing) of 18 and 24 individuals, respectively. In 2021, blocks contained six F_2_ hybrids and six of each parent, with Granite 2 having 75 blocks, totaling 4,050 plants. In 2022, blocks contained 16 F_3_ hybrids with four of each parent, totaling 4,800 plants. In 2023, we included local parental genotypes planted only at their origin site to compare fitness with the original parents (WLF and YVO). We used 2–3 intrapopulation parental crosses and original inbred lines, which did not differ in mean seed production and were therefore combined for subsequent analyses (Table S2). Each site had 60 blocks (5×7 formation) of 35 individuals, composed of 3**–**4 parental genotypes (YVO, WLF, and local), and 16**–**23 F_6_ hybrids per block, with 1,200 of each parent and 3,000 F_6_ hybrids per habitat totaling 8,400 plants. We used 300 F_6_ lines with 5 replicates per site.

We cold-stratified parental *M. laciniatus* and hybrid seeds for 10 days and parental *M. guttatus* seeds for 5 days at 4°C in the dark in plastic flats filled with soil (Sunshine Mix #4 Professional Growing Mix, Sun Gro) to break dormancy and synchronize germination for one week in greenhouses at the University of California (UC) Davis (2021) and UC Merced (2022**–**2023). Following germination upon the emergence of cotyledons, individuals were planted into block designs described above. Mortality within the first week was attributed to transplant shock, dead germinants were replaced and not included in subsequent analyses.

### Environmental & phenotypic data collection

To assess microhabitat variation, environmental measurements were taken weekly from experimental blocks. We measured soil moisture (%) using the SM150T sensor (Dynamax), soil surface temperature (°F) using a laser thermometer, and light intensity (µmol m-2 s-1) using a MQ-200X Sunlight Quantum Meter (Apogee Instruments). Light and temperature were not measured in 2013 (Ferris and Willis 2018). We recorded survival, herbivory, and putatively adaptive phenotypes under selection: flowering time, plant height, herkogamy, flower size, leaf shape (Ferris et al. 2014; Ferris and Willis 2018; Tataru et al. 2023). Foliar herbivory was assessed as the presence or absence of visible damage to leaf tissue. At first flowering, we recorded the date and measured plant height (mm), stigma and anther lengths (mm), and corolla width (mm). Flowering time was calculated as days from transplant to first flower. Herkogamy (i.e. stigma-anther separation) was calculated by subtracting stigma length from longest anther length, a metric of outcrossing ability. To assess leaf shape, the first true leaf was collected one-week post-flowering. Leaves were scanned and analyzed for leaf lobing index using ImageJ (Schneider et al. 2012; Ferris et al. 2015).

### Testing for effects of environmental factors on fitness

To identify environmental predictors of survival within each habitat, we pooled data across sites and used binomial mixed models with interval survival as the dependent variable, and soil moisture, soil temperature, light levels with their interactions as fixed independent variables, and block nested within site as a random effect. Interval survival was modeled as the number of survivors in each block at the start of a census period. We performed model selection for the best-fit model using AIC selection criteria using R packages *nlme* v3.1 (Pinheiro and Bates 2000; Pinheiro et al. 2023) and *MuMIn* v1.47.5 (Bartoń 2018). To test whether foliar herbivory differs between habitats, we used a binomial generalized linear mixed model with herbivory as the dependent variable and habitat as the fixed independent variable with block nested within site as a random effect. To understand the effect of herbivory on fitness, we fit Poisson generalized linear mixed models with fitness (i.e. seed count) as the dependent variable while herbivory and habitat and their interactions were fixed independent variables with block nested within site as a random effect using the R package *lme4* v1.1.32 (Bates et al. 2015).

### Phenotypic plasticity analyses

We used parental inbred lines to investigate each species’ phenotypically plastic response to the environmental variation between species’ habitats within individual years. These spatial contrasts provide a coarse-grained test (Levins 1968) of how genotypes may respond to differing environmental conditions. Plasticity was analyzed using generalized linear mixed models with phenotype as the dependent variable, habitat as a fixed effect and block nested within site as a random effect. Further, we examined phenotypic plasticity between habitats and years (2013-2023) using generalized linear mixed models with phenotype as the dependent variable, habitat and year with their interactions as fixed effects and block nested within site as a random effect. We also compared hybrid trait means between habitats which are not true measures of plasticity since they represent both genetic and environmental differences.

### Phenotypic selection analyses

Because *M. guttatus* and *M. laciniatus* are annual species, total reproductive output provides an estimate of lifetime fitness. We quantified reproductive output as fruit number (2013) or seed number (2019–2023). Plants that produced flowers but did not produce fruit or seed were assigned a value of zero. We calculated the correlation between fruit and seed number using Kendall’s correlation analysis. To facilitate comparisons to the 2013 experiment, we repeated all annual analyses using fruit number as the dependent variable. Selection gradients (/3 values) derived from fruit number were similar to those derived from seed number (Table S4), likely due to a positive correlation between seed and fruit number (Table S5). Therefore, we compared fruit and seed-based selection gradients across years.

To investigate linkage between traits, we calculated correlations between traits from hybrid individuals, grouped separately by habitat and year, using linear mixed models with traits as dependent and fixed variables with block nested within site as a random effect. Most correlations were weak (*r* < 0.50) and inconsistent across habitats and years, except between plant height and leaf area and between leaf area and flower width (Table S3). Therefore, leaf area was excluded from subsequent phenotypic selection analyses. Quantitative traits measured in each experiment (leaf lobing, plant height, outcrossing, flower width, flowering time) were standardized to a normal distribution with a mean of 0 and standard deviation of 1 to enable comparison of selection on qualitatively different traits and with previously published studies (Lande & Arnold 1983). For annual analyses, raw trait data were standardized within each year. For the combined analysis, raw trait data were combined across years and then standardized.

To understand annual fluctuations in the strength and direction of selection on traits in each habitat, we used datasets of hybrid individuals from each transplant year. We then used a dataset of all years combined to test for differences in whether spatial and temporal selection on a trait differed. To compare selection gradients from all years, data published in 2013 and 2019 were re-analyzed using the same multivariate linear selection model as the recent transplants (Ferris and Willis, 2018; Tataru *et al*. 2023). For annual phenotypic selection analyses, seed count (i.e. fecundity) was the dependent variable with plant height, flowering time, flower width, herkogamy, and leaf lobing as fixed independent variables, with block nested within site and habitat as a random effect. We did not include quadratic or correlational selection terms in our models to simplify our interpretation of the results (Anderson et al. 2015*b*). We did not conduct phenotypic selection analysis using an ASTER model approach because our dataset did not include trait data on individuals that died before flowering. To account for overdispersion of zeros in our fitness data we divided it into two components: 1) the combined probability of survival and reproduction (setting any seed) using binomial generalized linear mixed models and 2) the number of seeds set by plants that fruited (fecundity) using a zero-truncated Poisson generalized linear mixed model. Models were fit using the R package *glmmTMB* v1.1.8 (Brooks et al. 2017; Magnusson et al. 2017). Sites were pooled within habitats to give us the necessary statistical power.

To examine long-term selection, we repeated analyses on a combined dataset of hybrids pooled across all years (2013–2023) using fruit number as the dependent variable with plant height, flowering time, flower width, and leaf lobing as fixed independent variables, and block nested in site and year as random effects. Herkogamy was omitted due to its absence in 2013. To visualize the fitness-phenotype relationships, we plotted selection gradients using the R package *visreg* (Breheny and Burchett 2017). To assess whether slopes were different for traits between habitats, we used a bootstrap approach (1000 iterations) to derive 95% confidence intervals and *p*-values.

To assess whether temporally variation in phenotypic selection was related to environmental fluctuation, we used the combined dataset to fit generalized linear models for each habitat including interactions between traits and snowpack (snow water content accumulated by April 1^st^; Figure 1a). Models included fruit as the dependent variable, phenotypic traits and their interactions with snowpack as fixed independent variables and block nested within site and year as random effects. Lastly, we tested whether selection gradients differed among habitats and years using generalized linear models for each trait with habitat, year, and their interactions as fixed independent variables and block nested within site and habitat as a random effect.

### Assessing local adaptation of M. laciniatus vs. M. guttatus

To assess parental species’ local adaptation, we used three fitness components: survival to flowering (‘survival’), mean seeds per reproductive individual (‘fecundity’), and mean seeds per planted individual (‘total fitness’). For long term patterns, arithmetic means were calculated across years (2013, 2019, 2021, 2022, 2023) for each habitat and species. 2013 was omitted from averaged fecundity and total fitness because only fruit number was counted. Because it is not known whether they have seed banks, we also calculated geometric mean fitness across years. Genotype by environment (G×E) interactions were tested using linear mixed-effects models for fecundity and total fitness, and generalized linear mixed-effects models for survival, with parental genotype, habitat, and their interaction as fixed effects and block nested within site and habitat as a random effect. Finally, to assess the influence of the exceptionally high snowpack in 2023, we compared mean fitness with and without 2023. All models described above are summarized in Table S6.

## RESULTS

### Environmental variation causes spatial and temporal variation in survival

Fine-scale environmental variation was significantly associated with survival across habitats and years and fluctuated spatially and temporally with snowpack (Figure 2, Table S7). Soil moisture varied consistently between *M. laciniatus’s* granite and *M. guttatus’s* meadow habitat with moisture in granite outcrops declining rapidly after an early season plateau, while in meadows there is a gradual, linear decline throughout the growing season. Higher snowpack years (2019, 2023) produced shallower seasonal declines in soil moisture in both habitats (Figure 2). Light intensity and temperature were generally higher in granite, except in the exceptionally high snowpack year of 2023 when temperatures were cooler and similar between habitats (Figure 2; Table S7). Light intensity also varied seasonally, including a sharp decline in both habitats during August 2019, likely reflecting frequent thunderstorms that coincided with a temporary increase in soil moisture. Sites within a habitat generally exhibited similar patterns of environmental variation (Figure 2).

**Figure 2.**
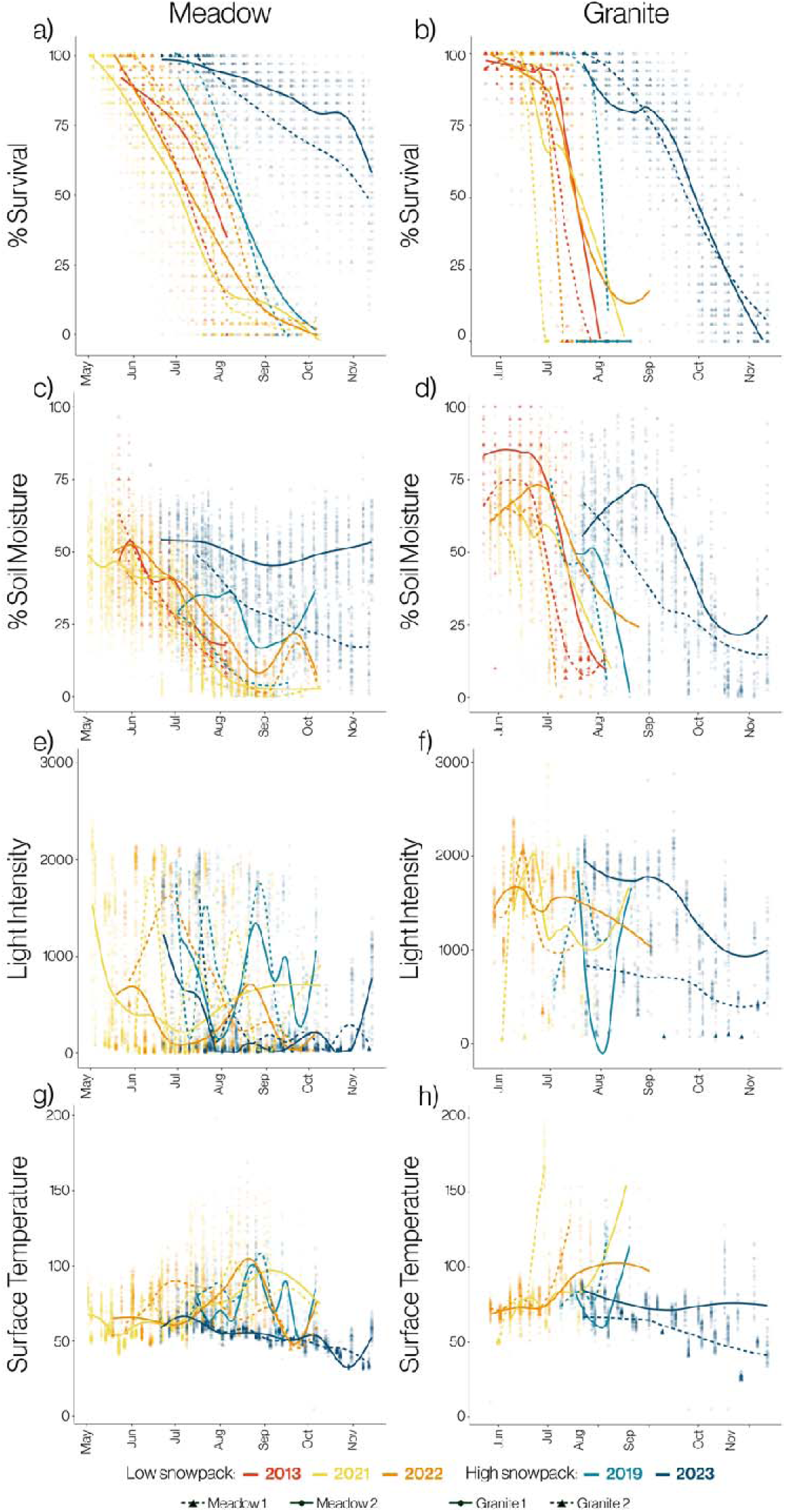
Seasonal changes in (a,b) survival (%), (c,d) soil moisture (%), (e,f) light intensity (µmol m-2 s-1), and (g,h) soil surface temperature (°F) in each block per site in meadow and granite habitats in low snowpack years (2013, 2021, 2022) and high snowpack years (2019, 2023). Lines represent fitted nonparametric smoothed curves. Sites were pooled together within habitats for further analyses. Data from 2013 and 2019 are published in Ferris and Willis 2018 and Tataru *et al*. 2023 and re-analyzed here.

Soil moisture was the most consistent predictor of interval survival across habitats and years, where it positively associated with survival in nearly all best-fit models (Table S7). The sole exception occurred in meadow habitat in 2023, where soil moisture was not significantly associated with survival (*p* > 0.05). This likely reflects the exceptionally high snowpack that year, which maintained high soil moisture throughout the growing season (Figure 2a). The effects of soil moisture were strongest during dry years (2013, 2021, and 2022) in both habitats, suggesting that water availability was a major determinant of survival in drought conditions. In contrast, the effects of light intensity and temperature were more variable among years and habitats, indicating that their influence on survival depended on environmental context. Consistent with this, many environmental variable interactions were significant in best-fit models in both habitats (Table S7). Overall, environmental predictor effects on survival were generally weaker and more variable in wet years than dry years, particularly in meadow habitats.

Similar to 2013 and 2019, herbivory pressure differed significantly between habitats with greater herbivory in *M. guttatus’*s meadows in 2021 (meadow 22%, granite 1.97%; *p* = 0.042), 2022 (meadow 37.75%, granite 2.58%; p < 0.01), and 2023 (meadow 23.17%, granite 8.74%; *p* = 0.032). In 2021 and 2023, herbivory and the interaction between herbivory and habitat significantly affected total fitness but did not in 2022. Generally, herbivore pressure varied spatially, while its effect on fitness varied temporally.

### Episodic selection: an extreme climatic event dramatically shifted the growing season

The extreme high snowpack year (2023) differed from other years in several aspects. The growing season was delayed by six weeks or more in both habitats, with surveys not beginning in granite until previous years’ surveys had ended (Figure 2a,b). Soil moisture declined more gradually in both habitats in 2023 than any previous year (Figure 2c,d), while light intensity and surface temperatures were lower (Figure 2e-h). In granite, there was no characteristic temperature spike in 2023 that usually coincided with mortality (Figure 2h). Unlike previous years, many plants had not senesced in meadows at the end of the growing season: 41.7% (Meadow 1) and 50.7% (Meadow 2) alive at the last survey before snowfall (Figure 2a). Of the individuals remaining in Meadow 1, many had not flowered indicating a significant life-history delay (0% local genotype, 31.0% hybrids, 44.1% *M. guttatus*, 70.7% *M. laciniatus*). Meadow 2 was similar (0% local genotype, 47.3% hybrids, 40.4% *M. guttatus*) except most individuals of *M. laciniatus* flowered (92.0%). The 2023 growing season was 36-70 days longer than previous years in granite habitats (Granite 2013-2022: 33-87 days; 2023: 112-113 days) but comparable to other years in meadows (Meadow 2013-2022: 74-160 days; 2023: 119-146 days).

### Phenotypes are plastic between habitats

Parental genotypes exhibited consistent phenotypic plasticity between habitats within years, with the non-native species generally shifting toward the phenotype of the local species. In all three years, *M. laciniatus* grew more *M. guttatus*-like in meadow habitat. Relative to granite habitat, plants flowered 4–10 days later, were 8–38 cm taller and produced larger leaves (196–8,911 mm² increase in leaf area; Tables S8, S9, S10). This is similar to patterns from 2013 except for flower size (Ferris & Willis 2018). In its nonnative granite habitat, *M. guttatus* shifted toward *M. laciniatus*-like traits, flowering 8–10 days earlier, growing 42–54 cm shorter, and producing smaller leaves (1,618–38,696 mm² reduction in leaf area; Tables S8, S9, S10). Furthermore, flowering time, plant height, flower width, leaf area displayed significant habitat × year interactions (all *p* < 0.000), indicating plasticity changed over time, (Table S11).

### Interannual fluctuation versus long-term directional selection

We found temporal and spatial fluctuations in the strength and direction of selection on each quantitative trait (Tables 1, 2; Figure 3). The combined probability of survival and reproduction models revealed how phenotypes affected whether or not plants reproduced whereas the fecundity selection models revealed how selection acted among individuals that reproduced (number of seeds). Based on observed species differences, we predicted selection in *M. laciniatus’s* granite habitat for early flowering time, smaller flowers, increased leaf lobing, shorter plants, and decreased stigma-anther distance (herkogamy), with the inverse direction in *M. guttatus’s* meadow habitat (Figure 3a-e). Although fecundity selection yielded a greater number of significant selection gradients, the direction of selection on the probability of reproduction was generally more consistent across years and habitats (Figure 3). This pattern suggests that selection acting before reproduction is more temporally consistent.

**Figure 3.**
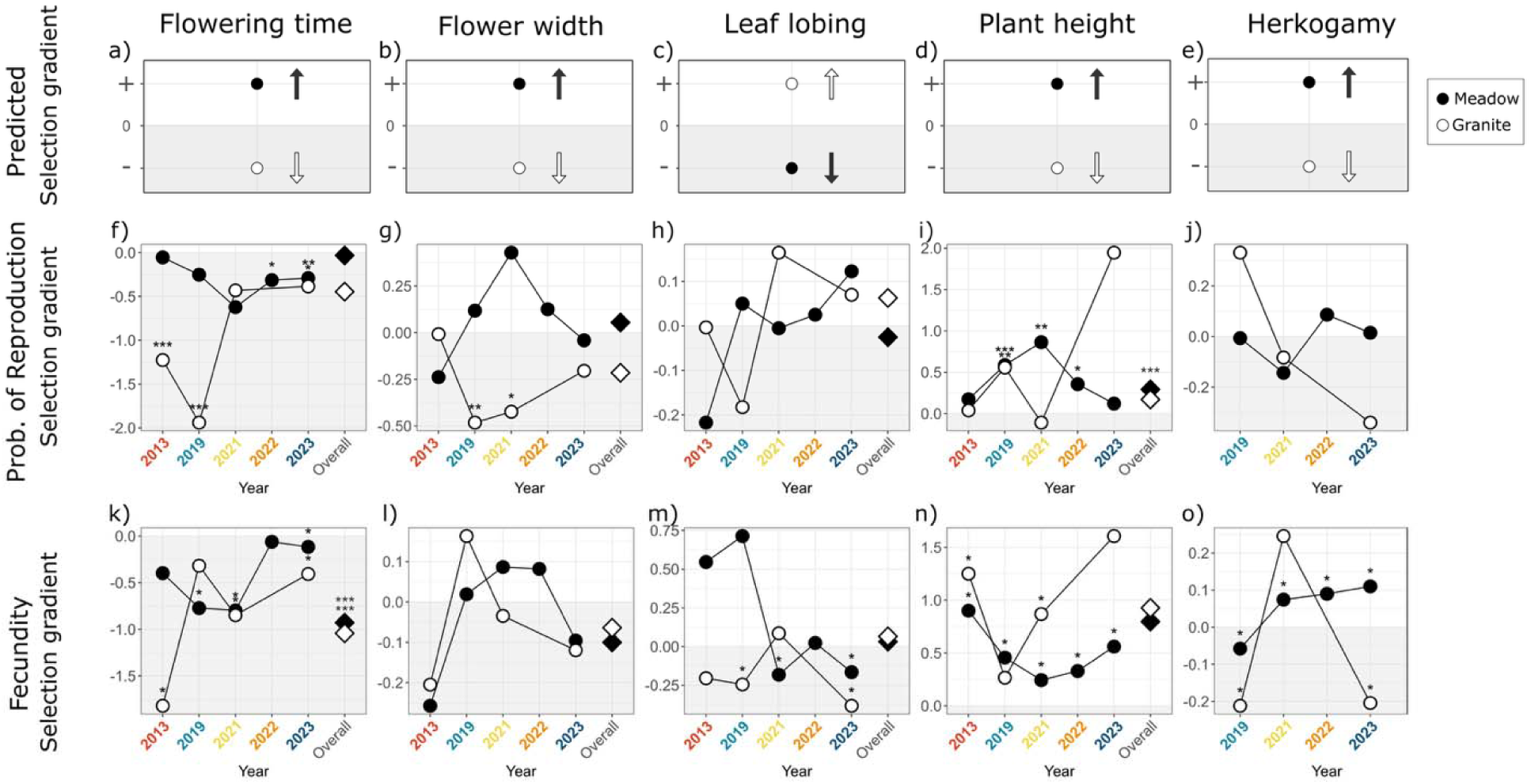
Visual representation of selection gradients (β) of phenotypic traits measured for hybrids during experimental years in granite (open circles) and meadow (filled circles) habitats and the overall estimate from all years combined (diamonds). Shown are (a-e) predicted values in each habitat based on native species trait differences and estimated values from (f-j) binomial models of selection on the combined probability of survival and reproduction and (k-o) zero-truncated Poisson models of fecundity selection. Significance of each β value is indicated with asterisks (* *p* < 0.05, ** *p* < 0.01, *** *p* < 0.001) and negative graph areas are shaded grey for clarity. All β values are based on seed number, except for values from 2013 which are based on fruit number, as seed number was not recorded. Herkogamy was excluded due to its absence in 2013. Data from 2013 and 2019 are published in Ferris and Willis 2018 and Tataru *et al*. 2023 and re-analyzed here.

**Table 1.**
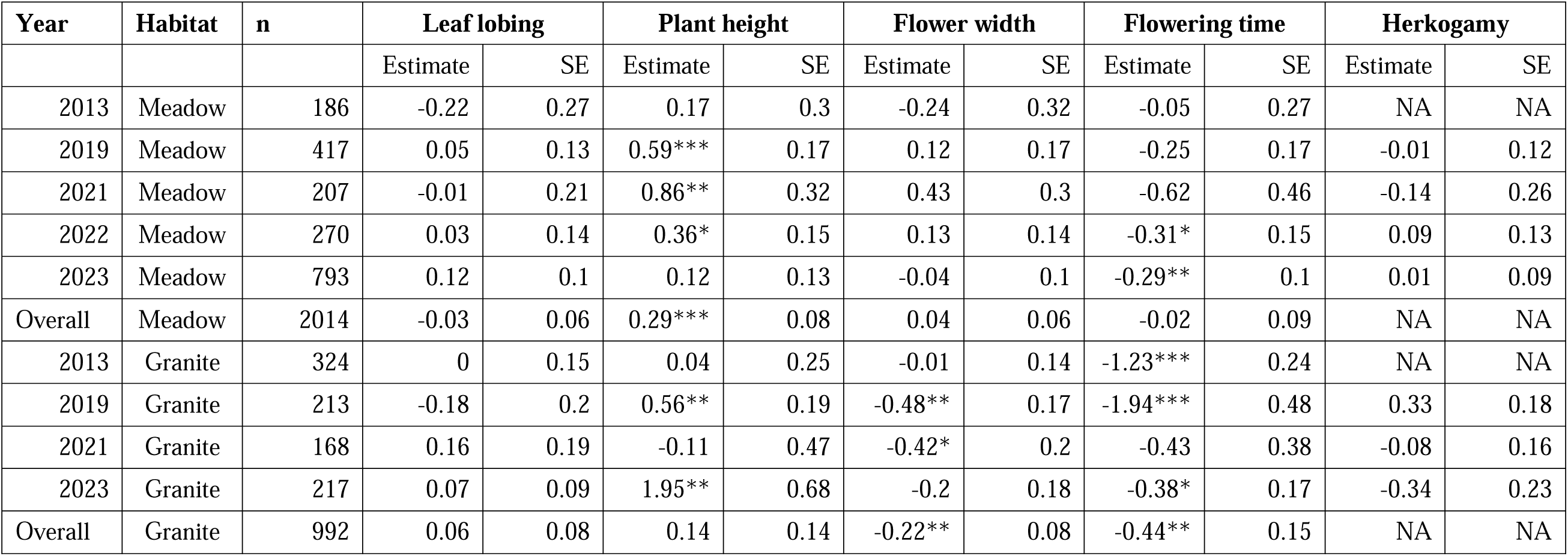
Analyses of the probability of survival and reproduction (binomial analysis) relating phenotypes to fitness, with fitness defined as whether or not a plant produced fruit (2013) or seeds (2019, 2021, 2022, 2023). A combined dataset of all years used fruit number for fitness. The strengths of selection for linear selection gradients are represented as β values. Asterisks indicate significance of selection in a trait: * *p* < 0.05, ** *p* < 0.01, *** *p* < 0.001. Herkogamy was excluded from the combined analysis due to its absence from the 2013 dataset. Phenotypic data from 2013 and 2019 are published in Ferris and Willis 2018 and Tataru *et al*. 2023 and re-analyzed here.

**Table 2.** Analyses of selection on fecundity (zero-truncated Poisson analysis) relating phenotypes to fitness, with fitness defined as the number of seeds produced, on individual experimental years of 2013, 2019, 2021, 2022, and 2023 and on a combined dataset of all years. The strengths of selection for linear selection gradients are represented as β values. Asterisks indicate significance of selection in a trait: * *p* < 0.05, ** *p* < 0.01, *** *p* < 0.001. Herkogamy was excluded to from the combined analysis due to its absence from the 2013 dataset. Phenotypic data from 2013 and 2019 are published in Ferris and Willis 2018 and Tataru *et al*. 2023 and re-analyzed here.

| Year | Habitat | n | Leaf lobing |  | Plant height |  | Flower width |  | Flowering time |  | Herkogamy |  |
| --- | --- | --- | --- | --- | --- | --- | --- | --- | --- | --- | --- | --- |
|  |  |  | Estimate | SE | Estimate | SE | Estimate | SE | Estimate | SE | Estimate | SE |
| 2013 | Meadow | 45 | 0.55 | 0.32 | 0.90* | 0.39 | -0.26 | 0.51 | -0.4 | 0.41 | NA | NA |
| 2019 | Meadow | 114 | 0.71*** | 0.04 | 0.46*** | 0.04 | 0.02 | 0.24 | -0.77*** | 0.06 | -0.06* | 0.02 |
| 2021 | Meadow | 129 | -0.18*** | 0.02 | 0.24*** | 0.02 | 0.09 | 0.11 | -0.80*** | 0.04 | 0.07*** | 0.02 |
| 2022 | Meadow | 139 | 0.02 | 0.02 | 0.33*** | 0.04 | 0.08 | 0.08 | -0.06* | 0.03 | 0.09** | 0.03 |
| 2023 | Meadow | 207 | -0.17*** | 0.02 | 0.56*** | 0.03 | -0.1 | 0.1 | -0.12*** | 0.03 | 0.11*** | 0.02 |
| Overall | Meadow | 547 | 0.04 | 0.04 | 0.79*** | 0.1 | -0.1 | 0.07 | -0.92*** | 0.13 | NA | NA |
| 2013 | Granite | 144 | -0.2 | 0.16 | 1.25*** | 0.28 | -0.2 | 0.17 | -1.82*** | 0.33 | NA | NA |
| 2019 | Granite | 66 | -0.24* | 0.1 | 0.27** | 0.09 | 0.16 | 0.12 | -0.32 | 0.23 | -0.21* | 0.09 |
| 2021 | Granite | 88 | 0.09 | 0.06 | 0.87*** | 0.15 | -0.03 | 0.15 | -0.85*** | 0.14 | 0.25*** | 0.05 |
| 2023 | Granite | 51 | -0.38*** | 0.09 | 1.61*** | 0.17 | -0.12 | 0.16 | -0.41*** | 0.06 | -0.20*** | 0.06 |
| Overall | Granite | 998 | 0.06 | 0.04 | 0.91*** | 0.08 | -0.06 | 0.05 | -1.02*** | 0.08 | NA | NA |

Flowering time showed the strongest and most consistent pattern of selection. In three of four years with comparable habitat estimates, there was stronger negative selection on flowering time in granite than meadow habitat both in terms of the probability of reproduction and in two years for fecundity selection. Differences in selection gradients between habitats ranged from 0.09 to 1.69 β units for the probability of survival and reproduction and from 0.05 to 1.42 units for fecundity selection, indicating stronger selection for early flowering in granite despite selection acting in the same direction in both habitats (Figure 3f,k). In contrast, patterns of selection on flower width were less consistent. The probability of survival and reproduction was consistently in the predicted direction, favoring larger flowers in meadow habitat and smaller flowers in granite habitat across in three out of four years (Figure 3g). Fecundity selection on flower width was more variable and matched this pattern in two of four years (Figure 3l). Selection on leaf lobing was variable across years. In low snowpack years, more lobed leaves tended to increase the probability of survival and reproduction in granite and decrease it in meadows, consistent with predicted species differences, although these estimates were not significant (Figure 3h). Fecundity selection on leaf lobing also fluctuated temporally and there was often selection for more lobed leaves in meadow, but not granite, contrary to predictions. Taller plants were generally favorable across habitats and years. Fecundity selection for greater plant-height was stronger in granite habitat in three of four years, with habitat differences ranging from 0.35 to 1.05 β units (Figure 3n), whereas effects on the probability of reproduction were inconsistent (Figure 3i). For herkogamy, fecundity selection generally matched predictions during high snowpack years (2019, 2023). In 2019, selection was stronger for shorter stigma-anther separation in granite than meadow, while in 2023 shorter stigma-anther separation was favored in granite and greater stigma-anther separation in meadow. Differences in selection gradients between habitats ranged from 0.15 to 0.31 β units in these years, whereas the pattern reversed in the low snowpack year 2021 (Figure 3o).

Temporal fluctuations in selection were associated with variation in annual snowpack (Table S12). In meadow, increasing snowpack weakened selection for taller plants in both analyses (both p < 0.001). Snowpack also altered leaf lobing’s effect on the probability of survival and reproduction, causing selection to shift from favoring less-lobed leaves in low snowpack years towards more lobed leaves in high snowpack years (*p* < 0.05). In granite habitat, increasing snowpack weakened fecundity selection for earlier flowering and larger flowers, as indicated by positive flowering-time × snowpack and negative flower-width × snowpack interactions (both p < 0.01; Table S12). Flowering time and plant height’s effects on the probability of reproduction were also impacted by snowpack in granite (p < 0.05; p < 0.001), where higher snowpack reduced selection for earlier flowering and increased selection for taller plants.

Fecundity selection on flowering time and plant height varied significantly with habitat and year (both *p* < 0.001; Tables S13, S14). In contrast, fecundity selection on flower width and leaf lobing did not fluctuate significantly over time (*p* > 0.05). The effect of flowering time (*p* < 0.05), flower width (*p* < 0.05), and plant height (*p* < 0.001) on the probability of survival and reproduction also varied significantly between habitats and years. Leaf lobing did not show evidence of variation in its effect on the probability of survival and reproduction (p > 0.05). Therefore, we find that temporal fluctuations in selection were strongest on flowering time and plant height.

Despite substantial interannual fluctuations combining our data across years revealed long-term patterns of directional selection (Table 1, Table 2; Figures 3, 4, 5). Regarding the probability of survival and reproduction, flowering time was under weak selection in meadow, whereas early flowering had a positive effect in granite following predictions (Figure 4a,b). Similarly, as predicted, smaller flower size had a positive effect in granite habitat, while larger flower size was under weak selection in meadows (Figure c,d). Leaf lobing increased the long-term probability of survival and reproduction in granite, while slightly decreasing it in meadows, following predictions (Figures 4e,f). Increased plant height increased the probability of reproduction in both habitats (Figure 4g,h). With respect to fecundity selection, we found long-term selection for earlier flowering time, smaller flowers, more lobed leaves and taller plants in both habitats (Figure 5). Bootstrap comparisons of selection gradients confirmed significantly stronger selection in granite for early flowering time (probability of reproduction: *p* < 0.0001), greater leaf lobing (fecundity: *p* < 0.0001), and larger plant sizes (fecundity: *p* < 0.0001). Therefore, while inter-annual fluctuations were common, long-term directional selection remained aligned with species’ differences in flowering time and leaf shape.

**Figure 4.**
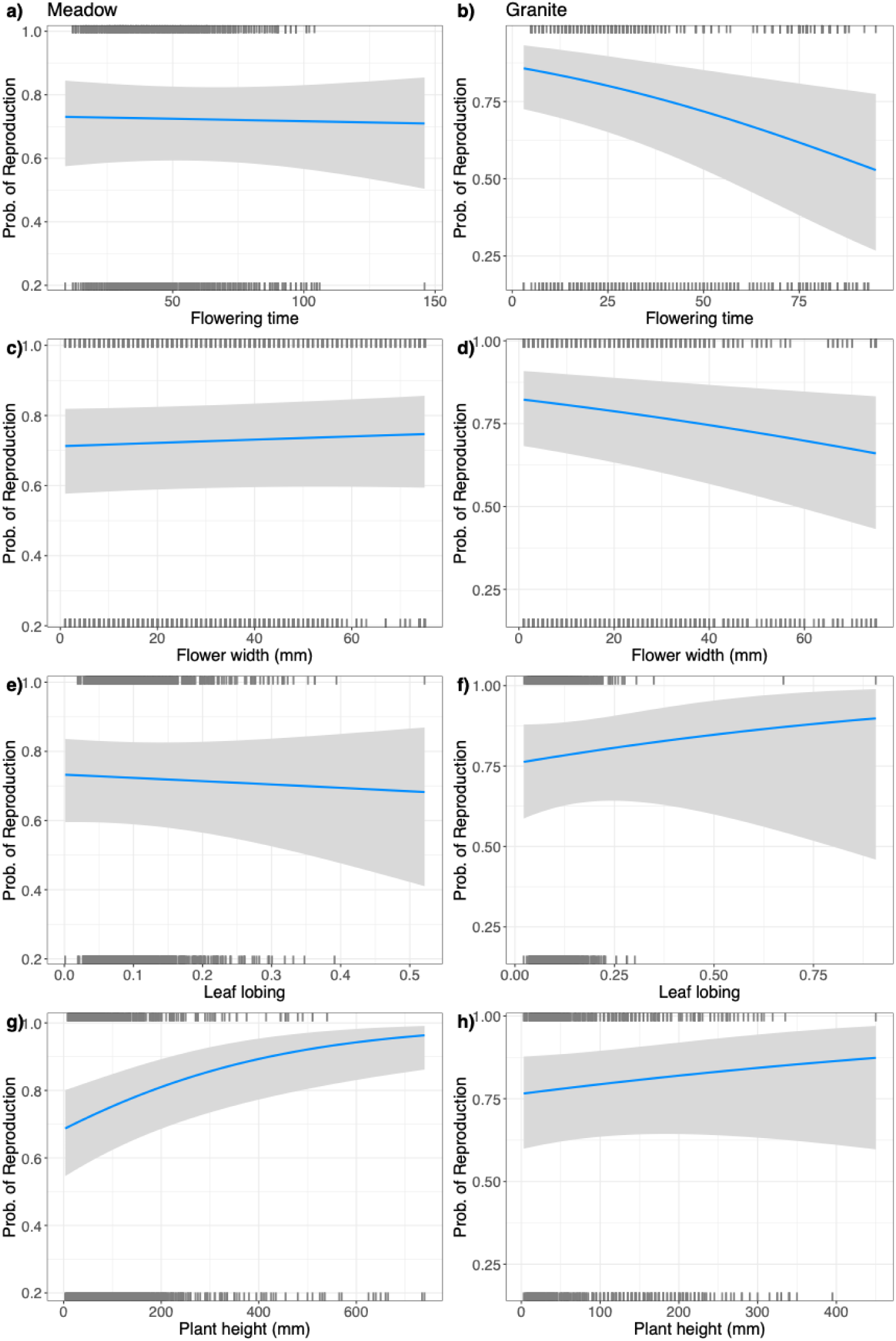
Predicted probability of reproduction with all years combined on meadow (left) and granite (right) on (a,b) flowering time, (c,d) flower width (mm), (e,f) leaf lobing index, and (g,h) plant height (mm). Curves show predictions from binomial generalized linear mixed models. Shaded regions are 95% confidence intervals. Marks along the x-axes indicate the distribution of observed trait values, which are unstandardized for biological relevance. Data from 2013 and 2019 are from Ferris and Willis 2018 and Tataru *et al*. 2023 and re-analyzed here.

**Figure 5.**
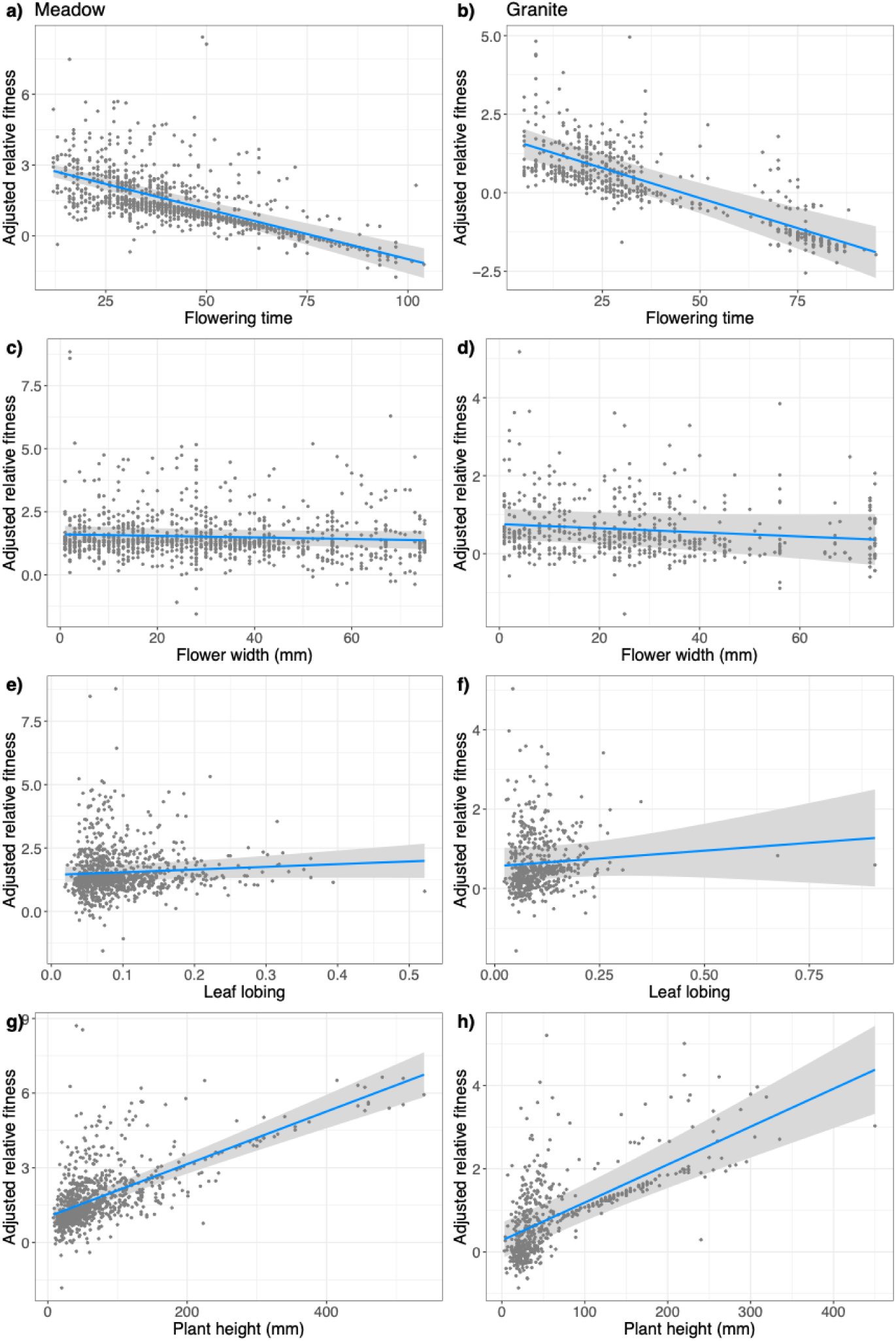
Cumulative fecundity selection with all years combined in meadow (left) and granite (right) habitat in a zero-truncated Poisson analysis on (a,b) flowering time, (c,d) flower width (mm), (e,f) leaf lobing index, and (g,h) plant height (mm). Y-axes show adjusted relative fitness (fruit number), statistically corrected for other variables included in the models, and the x-axes show unstandardized trait values for biological relevance. Shaded regions are 95% confidence intervals. Data from 2013 and 2019 are published in Ferris and Willis 2018 and Tataru *et al*. 2023 and re-analyzed here.

### Local adaptation fluctuates temporally with long-term maladaptation of M. guttatus

We found temporal fluctuation in patterns of local adaptation across years and fitness components. In 2013, a drought year, total fitness exhibited a clear pattern of local adaptation with each species achieving higher fitness in its native habitat (Figure 6). In other years reciprocal local adaptation was not evident, although local adaptation within individual habitats was observed (Figure 6m–q). For example, in all three drought years *M. laciniatus* had higher fitness in its native granite habitat than *M. guttatus* (Figure 6m,o,p), while *M. guttatus* was locally adapted in only two out of five years (Figure 6m,n).

**Figure 6.**
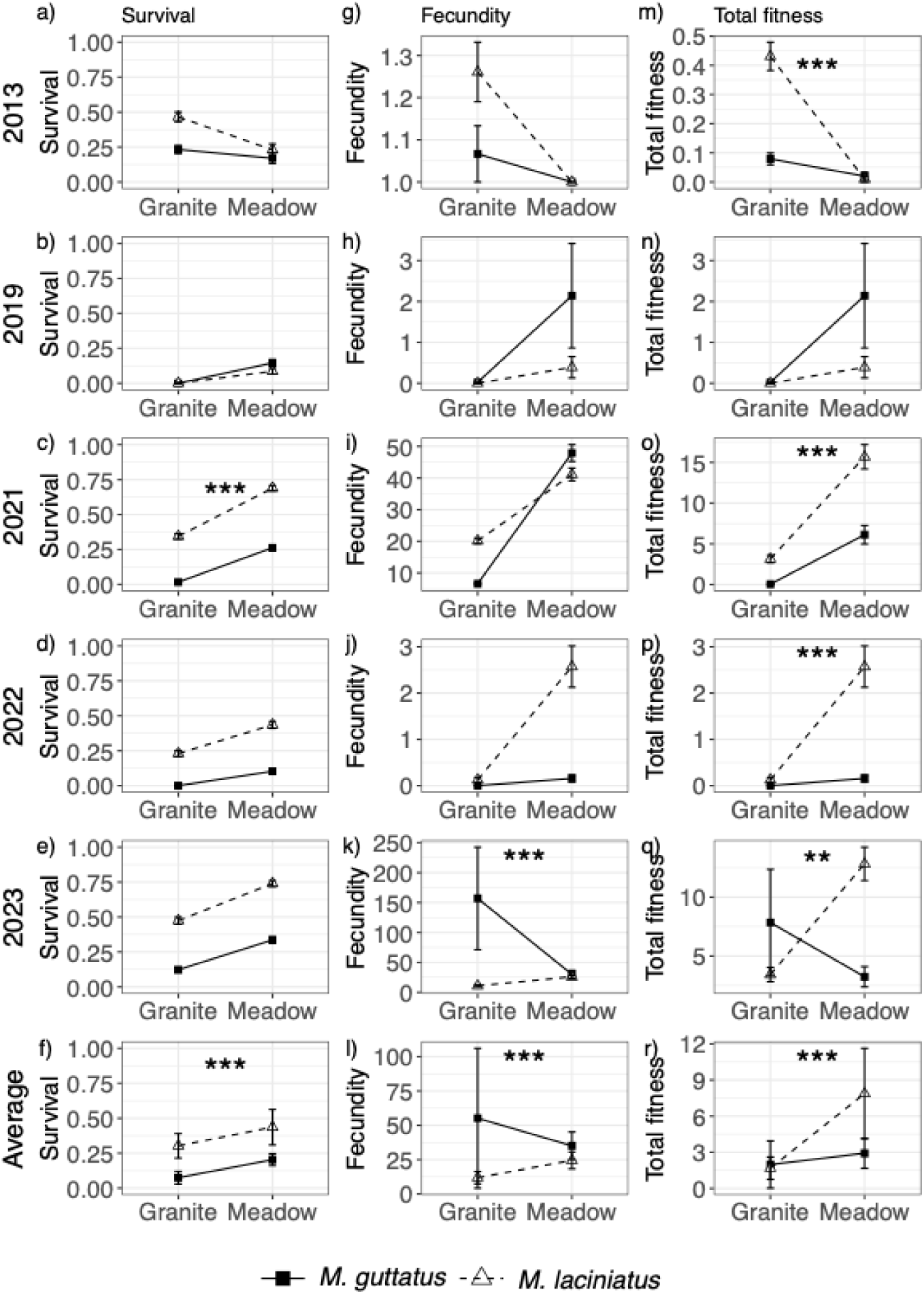
Survival to flowering (a-f), fecundity (mean seed number per reproductive individual; g-l), and total fitness (mean seed number per planted seed; m-r) of *Mimulus guttatus* (filled square, solid line) and *M. laciniatus* (open triangle, dashed line) in granite and meadow habitats, from each experimental year and the average (arithmetic mean) across years (f, l, r). Calculations of fecundity and total fitness for 2013 are based on fruit number because seed number was not recorded; 2013 was omitted from averaged fecundity and total fitness. Values from 2013 and 2019 are from Ferris and Willis 2018 and Tataru *et al*. 2023. Asterisks indicate significant G×E interactions: * *p* < 0.05, ** *p* < 0.01, *** *p* < 0.001.

Patterns of local adaptation also differed between the individual components of fitness: survival and fecundity. In each year *M. laciniatus* generally exhibited two to four times greater survival to flowering than *M. guttatus* in both habitats, with the exception of 2019 when survival was low for both species (Figure 6a-e) leading to higher average survival in both habitats (Figure 6f). Patterns of fecundity varied substantially between years. In 2013 and 2021, parental species achieved higher fecundity in their native habitats, consistent with local adaptation (Figure 6g-k). This pattern was strongest in 2021, when *M. laciniatus* produced three times more seeds than *M. guttatus* in granite and *M. guttatus* produced 20% more seeds than *M. laciniatus* in meadow (Figure 6i). In contrast, 2023 showed reciprocal maladaptation with *M. guttatus* producing ten times more seeds than *M. laciniatus* in granite, whereas *M. laciniatus* produced only slightly more seeds than *M. guttatus* in meadow (Figure 6k). The 2023 results strongly influenced the overall pattern of average fecundity with each species performing worse in its local environment than the foreign genotype (Figure 6l). However, excluding the exceptionally high-snowpack year of 2023 revealed a signature of reciprocal local adaptation (Figure S2).

Significant genotype by environment interactions (G×E) were detected for total fitness in every year except 2019 (Table S15), whereas we detected G×E in survival and fecundity in only 2021 and 2023, respectively (Table S15). For total fitness, low-snowpack years (2013, 2021, and 2022) were characterized by strong fitness advantages of *M. laciniatus*. While *M. laciniatus* was locally adapted to granite habitat in the low-snowpack years (Figure 6m,o,p), during high-snowpack years *M. laciniatus* and *M. guttatus* had similarly low total fitness in granite (2019, 2023; Figure 6n,q). In contrast, there was no clear pattern in local adaptation between high and low snowpack years in meadow habitat (Figure 6n,q). In the extreme high snowpack of 2023, where the growing season was delayed by over a month, there was reciprocal local maladaptation in total fitness which contributed to the signature of maladaptation in average total fitness (Figure S3). These outcomes suggest that snowpack may partially explain temporal variation in local adaptation.

Across years, *M. laciniatus* had higher total fitness in meadow habitat, whereas total fitness was low for both species in granite (Figure 6r, Figure S2). Geometric mean fitness analyses reinforced these patterns. Because a single year of very low fitness can strongly reduce long-term population growth, particularly in granite habitats lacking a persistent seed bank, geometric mean fitness provides a biologically relevant measure of long-term performance. Under this metric, *M. laciniatus* maintained higher geometric mean total fitness than the native *M. guttatus* in meadow habitat, whereas geometric mean fitness was effectively zero for both species in granite (Figure S3).

## DISCUSSION

We investigated spatial and temporal variation in selection on sympatric Monkeyflowers, *M. guttatus* and *M. laciniatus*, across replicated reciprocal transplants spanning low (2013, 2021, 2022) and high (2019, 2023) snowpack years, including one extreme year of episodic selection. We found that key environmental variables such as soil moisture, temperature, and light intensity fluctuated over space and time, impacting fitness and phenotypic plasticity. Selection on traits underlying reproductive isolation and habitat adaptation fluctuated annually, but long-term patterns in the direction and magnitude of selection aligned with species differences in flowering time. Strong, repeated selection favoring earlier flowering in granite habitats should maintain divergence in flowering phenology (Figure 4), a key component of temporal reproductive isolation. Local adaptation also fluctuated but on average, *M. guttatus* was maladapted to meadows habitats, indicating habitat isolation may be asymmetric and temporally dependent. Finally, we found that episodic selection, caused by an extreme snowfall event, disrupted local adaptation and eroded species boundaries.

### Inter-annual fluctuation in natural selection does not erase signatures of long-term divergence

Temporal and spatial variation in selection has been well studied for its role in maintaining genetic variation within species (Keep et al. 2021; Leidinger et al. 2021; Kelly 2022; Johnson et al. 2023). However, few studies have empirically tested how fluctuating selection influences the strength of ecological reproductive isolation between sympatric species (but see Campbell and Powers 2015). Here, we address this question by comparing the strength and direction of phenotypic selection in native *Mimulus* habitats and examining how these patterns changed across five repeated transplant experiments.

Although selection fluctuated substantially among years (Figure 3), flowering time exhibited the most consistent pattern of selection across habitats and years. Because flowering time is a key pre-zygotic isolating barrier in the *M. guttatus* species complex, repeated selection favoring earlier flowering in granite outcrops is likely to reinforce phenological and species divergence despite considerable temporal variation in selection (Lowry et al. 2008; Figures 4, 5). Previous studies have found that later-flowering *M. guttatus* generally achieve higher fecundity (Mojica and Kelly 2010; Mojica et al. 2012), especially in a longer, high snowpack growing season with lowered desiccation risk. Consistent with this, we found that that snowpack significantly influenced the strength of flowering-time selection in both habitats (Table S13). Although flowering time selection was negative in both habitats, the magnitude of selection for earlier flowering was often stronger in granite habitats (Figure 3k). While this does not represent divergent selection in the predicted direction, this difference in magnitude of flowering time selection could nevertheless reinforce species’ phenological divergence.

Contrary to our prediction that selection would favor smaller plant sizes in granite based on *M. laciniatus’s* short stature, larger plants consistently had higher fecundity in both habitats. This result is in line with broader evidence that body size at reproduction is a key determinant of fecundity across taxa (Aarseen and Taylor 1992; Aarssen and Jordan 2001; Kingsolver and Pfennig 2004). Nevertheless, average plant height remained smaller in granite than in meadow habitats (Table S9, Table S10, Table S11), likely reflecting ecological constraints imposed by limited water and nutrient availability that have led to the evolution of smaller size in rocky outcrops.

Selection on flower size fluctuated across years and habitats (Figure 3, Tables 1, 2). Selection on survival and reproduction favored smaller flowers in the granite habitat, consistent with *M. laciniatus* having smaller flowers. Across years, our combined analysis estimating trait’s effects on the probability of survival and reproduction revealed selection for smaller flowers in granite and weak selection for larger flowers in *M. guttatus’s* meadows (Figure 5, Table 1). Flower size affects reproductive success, is often associated with mating system differences, and can contribute to population differentiation and reproductive isolation (Bradshaw et al. 1995; Brunet 2009; Schiestl and Schluter 2009; Venail et al. 2010; Krizek and Anderson 2013). Flower size is often correlated with plant size indicating that early flowering, small stature, and small flowers are linked (Troth et al. 2018). Previous work in *M. guttatus* has found that flowering time alleles are pleiotropic, with later flowering alleles also increasing plant and flower size (Mojica et al. 2012; Monnahan and Kelly 2015). However, in our study these trait correlations changed in strength and direction among habitats and years (Table S2). Although larger flowers can increase fecundity, they may reduce viability in seasonally dry environments resulting in net selection for smaller flowers (Mojica and Kelly 2010). This supports our finding that selection favored earlier flowering and smaller flowers in *M. laciniatus’s* rapidly drying granite outcrops (Tables 1, 2).

Although individual yearly selection gradients for leaf lobing were not significant (Figure 3h), the relationship between leaf lobing and fitness varied predictably with snowpack across years, as indicated by a significant interaction with snowpack (Table S7). In low snowpack years, estimated selection gradients tended to favor more highly lobed leaves, whereas in high-snowpack years the direction of selection tended to reverse. Lobed leaves have a thinner boundary layer than round leaves of the same size which affects rates of heat and gas exchange as well as hydraulic resistance (Nicotra et al. 2011). Repeated evolution of lobed leaves in other members of the *M. guttatus* complex occupying rocky environments (Ferris et al. 2014) and altitudinal clines in leaf shape within *M. laciniatus* (Love and Ferris 2024) suggest that this trait is under environmentally mediated selection. Here we find that selection on leaf shape is temporally, as well as spatially, variable. Finally, we found little evidence for strong contemporary selection on herkogamy, suggesting that selection on stigma-anther separation is no longer as strong as during the earlier stages of speciation. This pattern is consistent with the expectation that strong selection on floral traits may be concentrated during episodes of diversification and weaken as reproductive barriers are more established (Harder and Johnson 2009).

By looking at patterns of selection in our combined dataset we found longer-term differences in the strength and direction of selection between these species’ habitats despite interannual fluctuations (Tables 1, 2; Figures 4, 5). Flowering time exhibited the clearest evidence of spatial variation in selection, with divergent selection between habitats in terms of the probability of reproduction (Table 1; Figure 4) and stronger fecundity selection for earlier flowering in granite than meadow (Table 2; Figure 5). Both types of selection favored earlier flowering in *M. laciniatus’s* granite outcrops, while in *M. guttatus’s* meadows, selection on flowering time was either near zero or in the same direction but weaker in magnitude, which is consistent with species differences in flowering phenology. Therefore, natural selection on flowering time likely contributes to the maintenance of temporal reproductive isolation, as has been observed in other *Mimulus* species (Ramsey et al 2003; Hall and Willis 2006; Lowry et al. 2008). Similar habitat-based differences in selection on flowering time have been documented in other systems; Ågren et al. (2017) found that selection on flowering time varied temporally in Italian and Swedish populations of *A. thaliana* but was consistently stronger in Italy. We also found weaker long-term divergent selection on leaf-shape, where the probability of survival and reproduction was largely unaffected by leaf shape in *M. guttatus’s* habitat, but leaf lobing increased the probability of reproduction in granite. Selection on leaf lobing varied predictably with snowpack across years (Table S13), and more highly lobed leaves tended to be favored during low-snowpack conditions. Given repeated evolutionary shifts in leaf shape within the *M. guttatus* species complex and the potential role of leaf morphology in adaptation to dry, exposed rocky environments (Nicotra et al. 2011; Ferris et al. 2015; Love and Ferris 2024), leaf shape may contribute to divergent environmental adaptation between species which would increase habitat isolation. These results indicate that while short-term fluctuating selection is not always in the direction of species differences, longer-term selection can still maintain species differences and ecological reproductive isolation. Particularly, repeated episodes of habitat-dependent selection on flowering time may contribute to the long-term maintenance of adaptive divergence despite substantial temporal variation in selection (Chapurlat et al. 2020).

### M. guttatus exhibits temporally variable local adaptation and long-term maladaptation

Organisms are expected to have a fitness advantage in their home environment because spatial heterogeneity should favor the evolution of local adaptation (Hastings 1983). However, temporal variation may constrain adaptation if there are opposing selection pressures over time (Stearns 1992). Therefore, we predicted that if habitat isolation is a reproductive barrier, each species would consistently outperform the other in its native habitat. Instead, local adaptation varied substantially among years (Figure 6m-q). Reciprocal local adaptation occured only in the drought year of 2013, whereas other years exhibited either local adaptation in one habitat or none at all. Across years, *M. laciniatus* maintained a higher average fitness than *M. guttatus* in both habitats, indicating long-term maladaptation of *M. guttatus*. This pattern mirrors our phenotypic selection analyses, which frequently favored *M. laciniatus*-like phenotypes, particularly earlier flowering, suggesting that contemporary selection pressures may have shifted toward phenotypes associated with the granite-adapted species.

Although local adaptation is thought to be widespread (Hereford 2009), reciprocal fitness trade-offs are not always found in transplant experiments (Bennett and Lenski 2007; Lowry et al. 2009). For example, in his classic meta-analysis Hereford (2009) found evidence of local adaptation, although the strength and consistency of trade-offs associated with adaptation of populations varied among studies. Most studies were conducted over limited spatial and temporal scales limiting conclusions of long-term costs of adaptation (Hereford 2009). Our results may therefore reflect relatively low fitness costs of granite adaptation, allowing *M. laciniatus* to perform well in both habitats (Fry, 1996; Leimu & Fischer, 2008). It is also possible that our *M. guttatus* lines were negatively affected by inbreeding depression, as a largely outcrossing species. However, this is unlikely as intrapopulation crosses and inbred lines performed similarly (Table S3). Alternatively, increasing drought frequency and climatic variability may have shifted environmental optima, generating an adaptive lag in *M. guttatus* if evolutionary responses have not kept pace with recent climate change (Lane et al. 2012; Mills et al. 2013; Kooyers et al. 2019). Similar patterns of adaptive lag and local maladaptation have been found in other populations of *M. guttatus* (Kooyers et al. 2019). If persistent, over time this could increase opportunities for colonization of meadow habitats by *M. laciniatus,* potentially increasing hybridization or lead to population decline of *M. guttatus*.

Asymmetry in local adaptation, or a lack of trade-offs, has been documented in other transplant experiments (Hereford 2009; Gosden et al. 2015; Latreille and Pichot 2017; Toll and Willis 2018). For example, in a repeated reciprocal transplant between serpentine and sandstone adapted populations of *Leptosiphon parviflorus,* Dittmar & Schemske (2023) found that serpentine populations had a temporally consistent local fitness advantage, whereas sandstone populations were locally adapted in only two out of four years. Similarly, we observed substantial temporal variation in local adaptation, including local adaptation of *M. guttatus* in meadow habitat in 2019 but local maladaptation in 2023 despite both years experiencing high snowpack.

Temporal fluctuations in the environment may favor increased phenotypic plasticity (Kawecki & Ebert 2004). Because local adaptation and plasticity can evolve simultaneously, populations experiencing greater environmental variability may evolve broader environmental tolerance. The average fitness advantage of *M. laciniatus* across environments could be due to adaptive plasticity, specifically adaptation to a history of environmental heterogeneity in the rocky outcrop habitat (Ghalambor et al. 2015). We found that both *M. guttatus* and *M. laciniatus* exhibit plasticity in the direction of the local species’ phenotype in each habitat (Tables S8, S9, S10). However, *M. guttatus* was more plastic than *M. laciniatus* for all measured phenotypes except fitness, suggesting that plasticity alone is unlikely to explain the fitness advantage of *M. laciniatus*. The success of *M. laciniatus* in meadow habitat may also reflect release from the environmental stresses that characterize granite outcrops, demonstrating plasticity in fitness.

### Episodic selection weakens species divergence

The high snowpack of 2023 delayed the onset of spring by 4–6 weeks, producing atypical environmental conditions that differed from other high-snowpack years (Figure 2). These conditions contributed to local maladaptation (Figure 6k), as signatures of local adaptation in fecundity reappeared when 2023 was excluded (Figure S2). In meadow habitat, delayed flowering reduced fitness in hybrids and *M*. *guttatus* and no local meadow genotypes flowered before snowfall. However, many individuals remained alive at the first snowfall indicating potential facultative perenniality. In contrast, *M. laciniatus* was less affected, likely because it flowers rapidly under shorter daylengths (Friedman and Willis 2013; Ferris and Willis 2018; Love and Ferris 2024). Delayed flowering may have resulted from shorter day lengths failing to meet critical photoperiod thresholds (Friedman and Willis 2013; Fishman et al. 2014; Kenney and Sweigart 2016), weaker end-of-season drought cues (Kooyers et al. 2015; Mantel and Sweigart 2019), and low light (Figure 2).

In granite habitats, the delayed snowmelt fostered unusually favorable environments for the non-native *M. guttatus* leading to higher survival (12% of total planted; 24 individuals) compared to previous years (0-2% of total planted) and remarkably high fecundity (Figure 6e,k) exceeding *M. laciniatus* in its native habitat for the first time (Figure 6q). Despite the delayed season, most plants completed reproduction before snowfall in granite, supporting the importance of drought stress and seasonal moisture dynamics in regulating flowering phenology. High-elevation populations of *M. laciniatus* may also possess increased plasticity in critical photoperiod (Love and Ferris 2024).

Our results demonstrate that extreme climatic events may disrupt species boundaries by disrupting local adaptation and altering patterns of selection. Over longer timescales, episodic selection may shift parental ranges, promote hybridization or a new hybrid species (Grant and Grant 1993), or alter adaptive potential (Campbell-Staton et al. 2017). More longitudinal field studies are needed to understand long-term evolutionary consequences of these rare and unpredictable climatic events (Anderson 2016; Bailey and van de Pol 2016). For instance, do these events drive evolutionary change and does this outweigh selection acting during ‘normal’ periods? For species with limited ranges, such as the Sierra Nevada endemic *M. laciniatus*, will these events influence extinction risk of populations?

### Conclusions

Our repeated reciprocal transplants reveal how fluctuating selection influences species’ boundaries. The foreign advantage of *M. laciniatus* in the non-native meadow habitat across years suggests either low cost of adaptation to the granite habitat or recent maladaptation of *M. guttatus* due to environmental shifts. Furthermore, longer-term divergent selection on flowering time should reinforce temporal and habitat isolation between these species. However, the lack of consistent divergent selection on other key traits could lead to eventual species fusion. Finally, a bout of episodic natural selection due to extreme snowfall shifted population dynamics with potential long-term impacts on species’ range, abundance, and gene flow. Our study illustrates the strength of integrating field experiments across multiple spatial and temporal scales to gain a deeper understanding of the maintenance of biodiversity under environmental change (Wadgymar et al. 2017; Dittmar and Schemske 2023; Oakley et al. 2023).

## DATA AVAILABILITY STATEMENT

The data that support the findings of this study will be openly available in [repository name] at http://doi.org/[doi], reference number [reference number], upon acceptance of the manuscript.

## Supplementary Information

**Figure S1.**
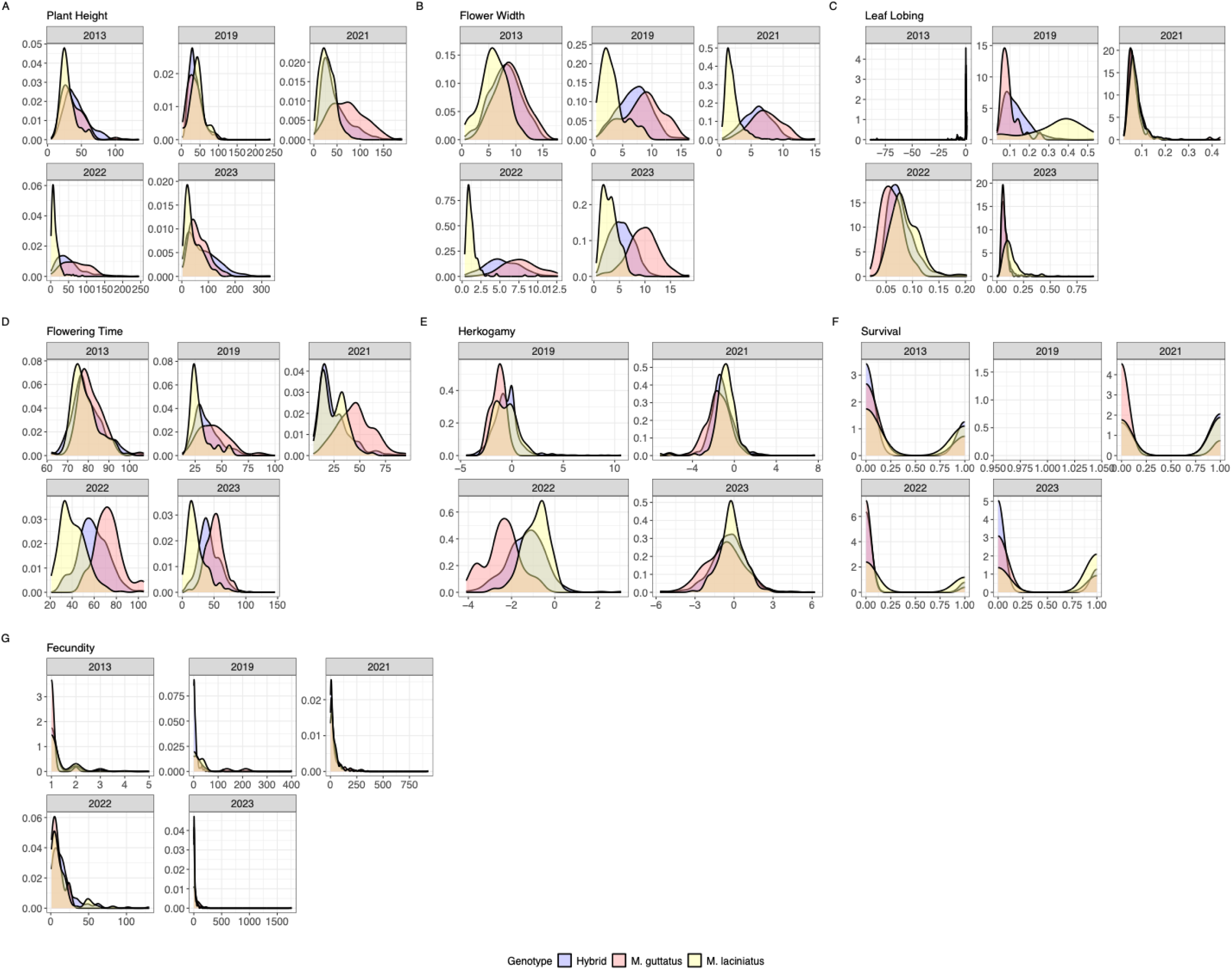
Density histograms of hybrid and parental values for phenotypic traits a) flower width, b) flowering time, c) plant height, d) leaf lobing, e) herkogamy, f) survival, and g) fecundity.in study years (2013, 2019, 2021, 2022, 2023). Herkogamy was not measured in 2013.

**Figure S2.**
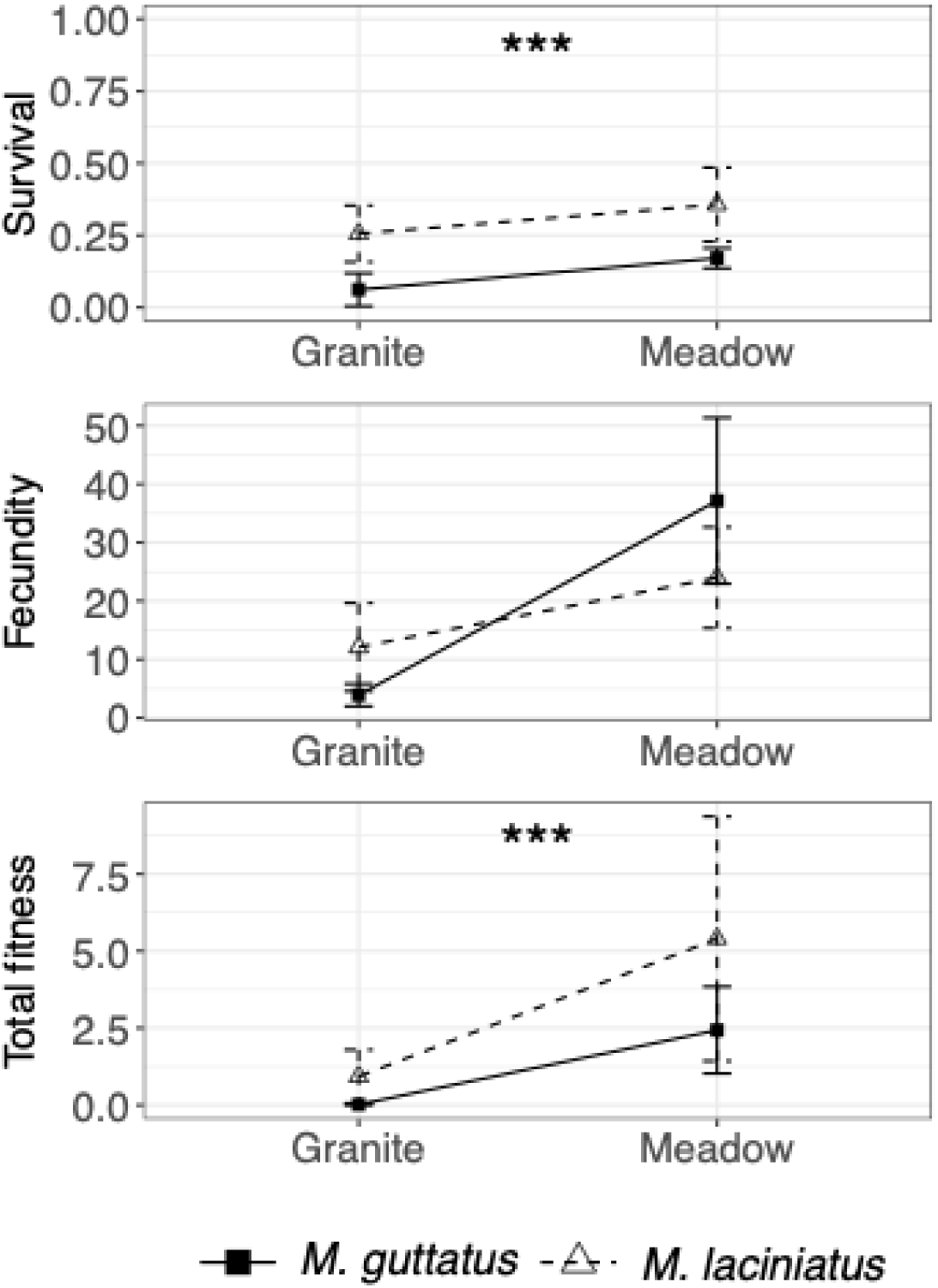
The average (arithmetic mean) of fitness metrics excluding 2023: survival to flowering, fecundity (mean seed number per reproductive individual), and total fitness (mean seed number per planted seed) from 2013-2022 of *Mimulus guttatus* (filled square, solid line) and *M. laciniatus* (open triangle, dashed line) in granite and meadow habitats. 2013 was omitted from fecundity and total fitness calculations because seed number was not recorded, only fruit number. Values from 2013 and 2019 are from Ferris and Willis 2018 and Tataru *et al*. 2023. This corresponds to Figure 5f, l, r.

**Figure S3.**
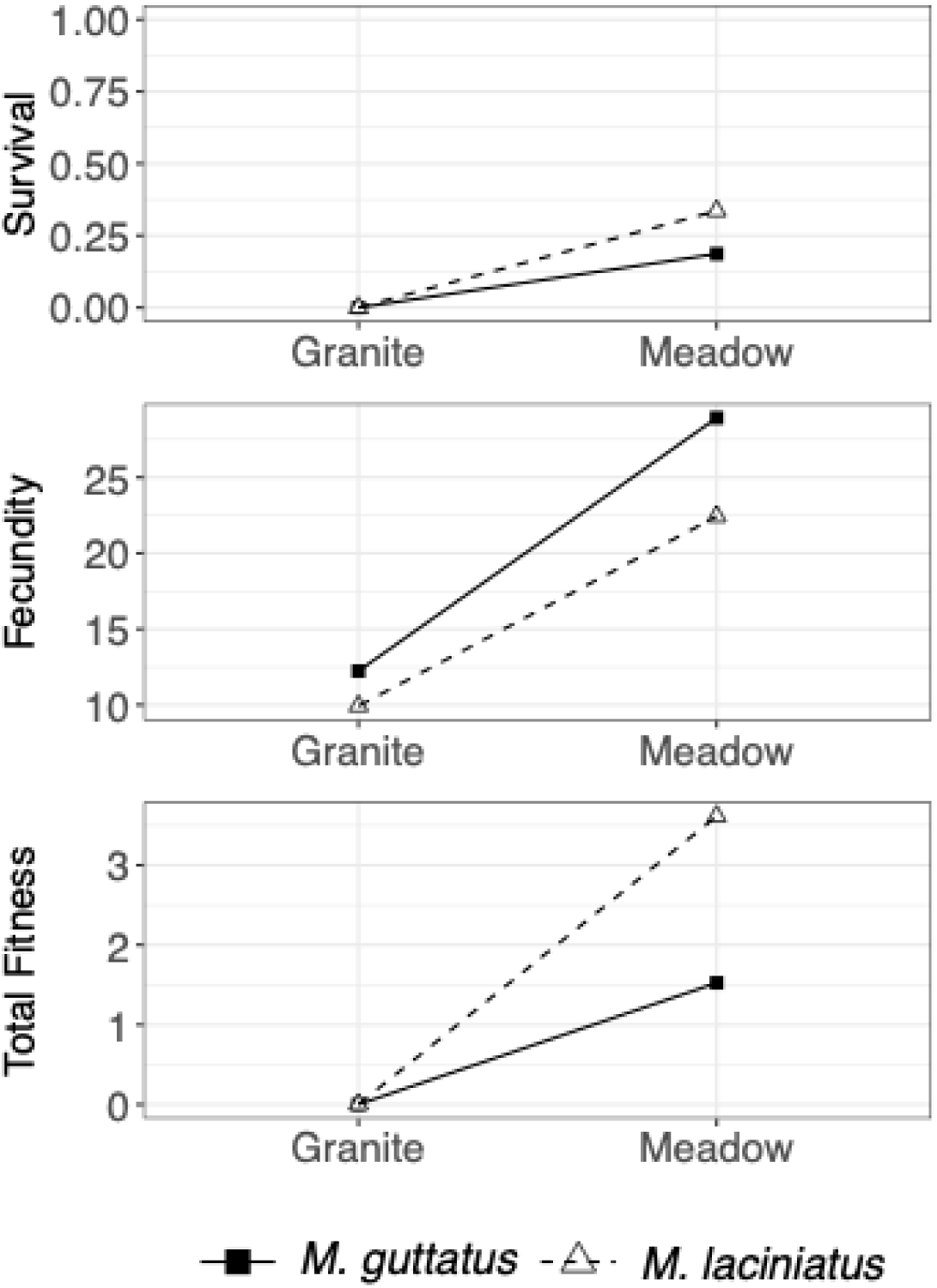
The average (geometric mean) of combined survival to flowering, fecundity (mean seed number per reproductive individual), and total fitness (mean seed number per planted seed) of *Mimulus guttatus* (filled square, solid line) and *M. laciniatus* (open triangle, dashed line) in granite and meadow habitat from experimental years. These values represent geometric means. 2013 was omitted from fecundity and total fitness calculations because seed number was not recorded, only fruit number. Data from 2013 and 2019 are from Ferris and Willis 2018 and Tataru *et al*. 2023 and re-analyzed here.

**Table S1.** Hybrid genotypes and generations used each experimental year, with sample sizes of each genotype planted at each site with total experimental sample sizes. The maternal genotype (WLF 47) is *M. laciniatus*, and the paternal genotypes (YVO 6 and 18) are *M. guttatus*. Details of the 2013 and 2019 experiments are published in Ferris and Willis 2018 and Tataru *et al*. 2023.

| Year | Hybrid genotype | Granite 1 | Granite 2 | Meadow 1 | Meadow 2 | Total |
| --- | --- | --- | --- | --- | --- | --- |
| 2013 | F4 WLF 47<br>maternal x YVO 6<br>paternal | 750 hybrids,<br>100 maternal,<br>100 paternal | 750 hybrids,<br>100 maternal,<br>100 paternal | 750 hybrids,<br>100 maternal,<br>100 paternal | 750 hybrids,<br>100 maternal,<br>100 paternal | 3,800 |
| 2019 | F4 WLF 47<br>maternal x YVO 6<br>paternal | 600 hybrids,<br>60 maternal,<br>60 paternal | 660 hybrids,<br>44 maternal,<br>88 paternal | 750 hybrids,<br>71 maternal,<br>100 paternal | 750 hybrids,<br>50 maternal,<br>100 paternal | 3,330 |
| 2021 | F2 WLF 47<br>maternal x YVO<br>18 paternal | 750 hybrids,<br>750 maternal,<br>750 paternal | 750 hybrids,<br>750 maternal,<br>750 paternal | 600 hybrids,<br>600 maternal,<br>600 paternal | 600 hybrids,<br>600 maternal,<br>600 paternal | 4,050 |
| 2022 | F3 WLF 47<br>maternal x YVO<br>18 paternal | 800 hybrids,<br>200 maternal,<br>200 paternal | 800 hybrids,<br>200 maternal,<br>200 paternal | 800 hybrids,<br>200 maternal,<br>200 paternal | 800 hybrids,<br>200 maternal,<br>200 paternal | 4,800 |
| 2023 | F6 WLF 47<br>maternal x YVO<br>18 paternal | 1500 hybrids,<br>200 maternal,<br>200 paternal,<br>200 local parent | 1500 hybrids,<br>200 maternal,<br>200 paternal,<br>200 local parent | 1500 hybrids,<br>200 maternal,<br>200 paternal,<br>200 local parent | 1500 hybrids,<br>200 maternal,<br>200 paternal,<br>200 local parent | 8,400 |

**Table S2.** Intrapopulation parental crosses used in the 2023 experiment. Due to insufficient seed stock of single crosses of YVO and WLF, we used 2-3 crosses and original inbred lines, distributed evenly across blocks and sites. With habitats combined, WLF genotypes did not differ significantly in total seed count (F = 2.324, df = 2, p = 0.099). Similarly, YVO genotypes also did not differ significantly (F = 1.944, df = 3, p = 0.124).

| Habitat | Genotype | Survival to flowering | %Survival | Seed count | Mean Seed Count | Seed SD | N |
| --- | --- | --- | --- | --- | --- | --- | --- |
| Granite | WLF60xWLF47 | 81 | 44 | 585 | 7.31 | 21.27 | 184 |
|  | WLF61xWLF47 | 108 | 66 | 784 | 7.26 | 12.78 | 164 |
|  | WLF47 inbred | 0 | 0 | 0 | NA | NA | 51 |
|  |  |  |  |  |  | **Total:** | **399** |
|  | YVO18xYVO26 | 0 | 0 | 0 | NA | NA | 20 |
|  | YVO25xYVO18 | 12 | 8 | 417 | 32.08 | 67.69 | 159 |
|  | YVO26xYVO18 | 35 | 18 | 961 | 27.46 | 51.83 | 200 |
|  | YVO18 inbred | 0 | 0 | 0 | NA | NA | 20 |
|  |  |  |  |  |  | **Total:** | **399** |
| Meadow | WLF60xWLF47 | 97 | 58 | 1760 | 17.25 | 32.41 | 166 |
|  | WLF61xWLF47 | 156 | 83 | 3201 | 20.13 | 33.95 | 187 |
|  | WLF47inbred | 35 | 76 | 156 | 4.46 | 8.43 | 46 |
|  |  |  |  |  |  | **Total:** | **399** |
|  | YVO18xYVO26 | 26 | 47 | 208 | 8 | 16.78 | 55 |
|  | YVO25xYVO18 | 26 | 23 | 378 | 14.54 | 47.43 | 114 |
|  | YVO26xYVO18 | 55 | 32 | 687 | 12.27 | 27.66 | 171 |
|  | YVO18 inbred | 26 | 44 | 21 | 0.81 | 2.77 | 59 |
|  |  |  |  |  |  | **Total:** | **399** |

**Table S3.**
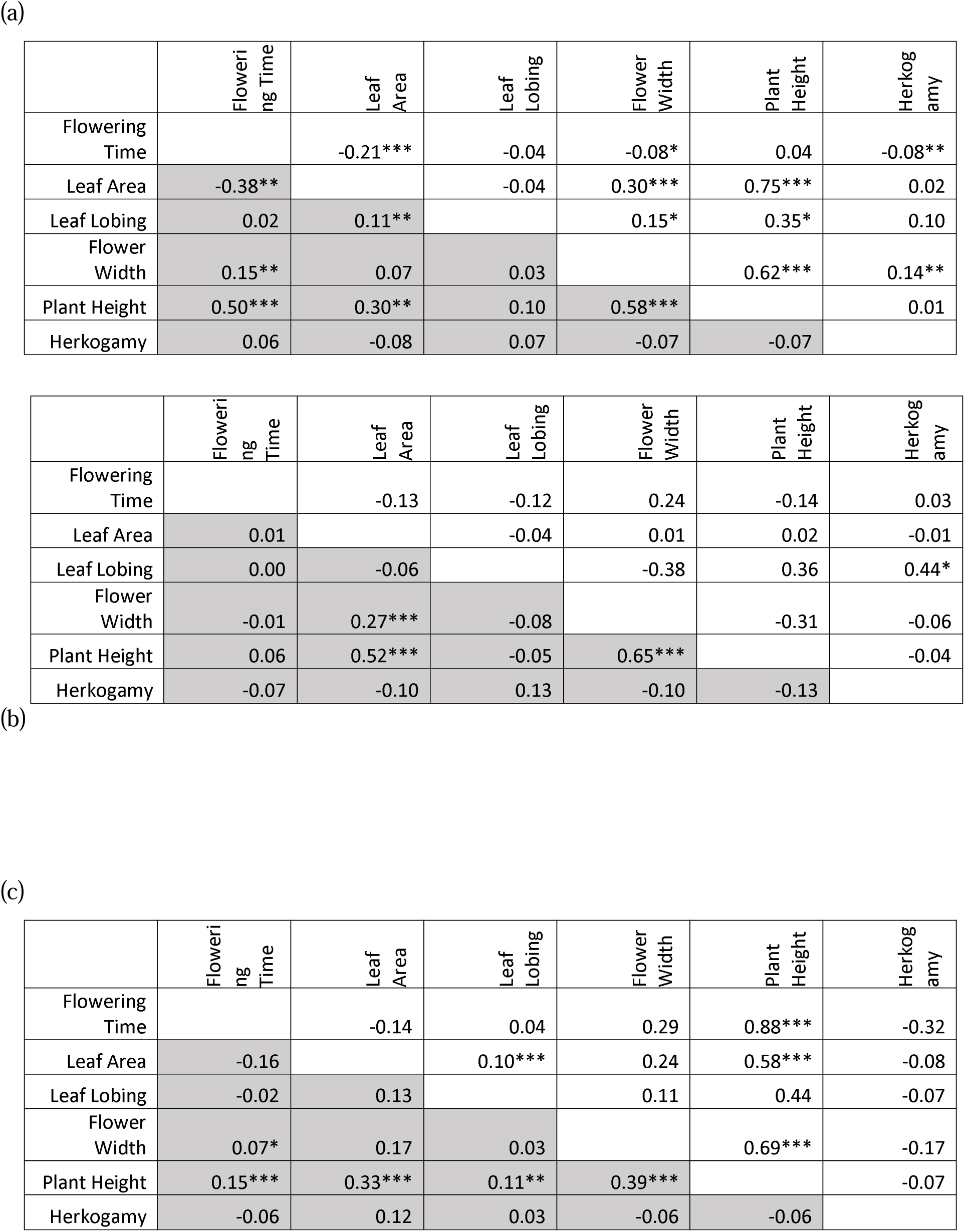
Hybrid phenotypic trait correlations in meadow (grey shaded; lower triangle) and granite (upper triangle) habitats for a) 2021, b) 2022, and c) 2023. Asterisks indicate significant (* = p < 0.05, ** = p < 0.01. *** = p < 0.001).

**Table S4.** Zero-truncated Poisson analysis (fecundity selection) on number of fruit produced if any, and binomial analysis (selection of survival and reproduction) on whether or not a plant produced fruits on individual experimental years of 2019, 2021, 2022, and 2023. The strengths of selection for linear selection gradients are represented as β values. Asterisks indicate significance of selection in a trait: ***p < 0.001; **p < 0.01; *p < 0.05. Data from 2019 are published in Tataru *et al*. 2023 and re-analyzed here.

| Selection | Year | Habitat | Leaf lobing |  | Plant height |  | Flower width |  | Flowering time |  |
| --- | --- | --- | --- | --- | --- | --- | --- | --- | --- | --- |
|  |  |  | Estimate | SE | Estimate | SE | Estimate | SE | Estimate | SE |
| Fecundity | 2019 | Granite | 0.14 | 0.07 | 0.17*** | 0.05 | -0.07 | 0.08 | -0.65*** | 0.18 |
| Fecundity | 2021 | Granite | 0.01 | 0.11 | 1.33*** | 0.33 | 0.11 | 0.14 | -0.82** | 0.27 |
| Fecundity | 2023 | Granite | -0.12 | 0.14 | 2.68*** | 0.59 | -0.40* | 0.17 | -0.40* | 0.19 |
| Fecundity | 2019 | Meadow | 0.12 | 0.07 | 0.42*** | 0.07 | -0.07 | 0.14 | -0.55*** | 0.12 |
| Fecundity | 2021 | Meadow | -0.01 | 0.06 | 0.43*** | 0.05 | -0.02 | 0.07 | -0.80*** | 0.11 |
| Fecundity | 2022 | Meadow | 0.13 | 0.12 | 0.52*** | 0.14 | 0.06 | 0.21 | -0.15 | 0.17 |
| Fecundity | 2023 | Meadow | 0.05 | 0.06 | 0.33** | 0.12 | -0.26* | 0.11 | -0.69*** | 0.11 |
| Prob. of Reproduction | 2019 | Granite | -0.25 | 0.22 | 0.25 | 0.25 | -0.32 | 0.23 | -0.53 | 0.56 |
| Prob. of Reproduction | 2021 | Granite | 0.23 | 0.21 | -0.21 | 0.46 | -0.50* | 0.2 | -0.3 | 0.37 |
| Prob. of Reproduction | 2023 | Granite | 0.18 | 0.15 | 2.46*** | 0.69 | 0.11 | 0.17 | -0.18 | 0.14 |
| Prob. of Reproduction | 2019 | Meadow | -0.09 | 0.12 | 0.14 | 0.16 | 0.13 | 0.18 | -0.19 | 0.15 |
| Prob. of Reproduction | 2021 | Meadow | -0.16 | 0.22 | 0.88* | 0.35 | 0.39 | 0.31 | -0.41 | 0.49 |
| Prob. of Reproduction | 2022 | Meadow | 0.04 | 0.15 | 0.21 | 0.16 | 0.17 | 0.15 | -0.18 | 0.15 |
| Prob. of Reproduction | 2023 | Meadow | 0.16 | 0.1 | 0.12 | 0.11 | 0 | 0.09 | 0.02 | 0.09 |

**Table S5.** Strength of correlations between fruit number and seed number calculated using Kendall’s correlation analysis.

| **Year** | **Habitat** | **$\tau$** | **p** | **n** |
| --- | --- | --- | --- | --- |
| 2021 | Meadow | 0.56 | 0.001 | 1296 |
|  | Granite | 0.59 | 0.001 | 809 |
| 2022 | Meadow | 0.64 | 0.001 | 545 |
| 2023 | Meadow | 0.75 | 0.0001 | 1393 |
|  | Granite | 0.69 | 0.0001 | 480 |

**Table S6.** Summary of models described in methods.

| Description of analysis | Model | Reference |
| --- | --- | --- |
| environmental variables and interval survival | glmmTMB(Interval survival ~ Soil moisture + Light + Temperature + Soil moisture × Light + Soil moisture × Temperature + Light × Temperature + Soil moisture × Light × Temperature + (1 \| Site/Block), family = binomial) | Figure 2, Table S7 |
| annual phenotypic (2019-2023), with family=binomial or truncated_poisson | glmmTMB(Seed ~ Leaf lobing + Plant height + Flower width + Flowering time + Herkogamy + (1 \| Site/Block)) | Figure 3, Tables 1 and 2 |
| annual phenotypic (2013), with family=binomial or truncated_poisson | glmmTMB(Fruit ~ Leaf lobing + Plant height + Flower width + Flowering time + (1 \| Site/Block)) | Figure 3, Tables 1 and 2 |
| phenotypic (years combined), survival/reproduction | glmmTMB(Fruit ~ Flowering time + Flower width + Leaf lobing + Plant height + (1 \| Site/Block) + (1 \| Year)), family=binomial | Figure 3f-i (overall diamonds), Figure 5 |
| phenotypic (years combined), fecundity | glmmTMB(Fruit ~ Flowering time + Flower width + Leaf lobing + Plant height + (1 \| Site/Block) + (1 \| Year)), family=truncated_poisson | Figure 3k-n (overall diamonds), Figure 4 |
| G×E (survival) | glmmTMB(Survival_Reproduction ~ Plant * Habitat + (1 \| Habitat/Site/Block), family = binomial(link = "logit")) | Figure 6a-f, Table S15a |
| G×E (fecundity) | glmmTMB(Fecundity ~ Genotype + Habitat + Genotype × Habitat + (1 \| Habitat/Site/Block)) | Figure 6g-l, Table S15b |
| G×E (total fitness) | glmmTMB(Total Fitness ~ Genotype + Habitat + Genotype × Habitat + (1 \| Habitat/Site/Block)) | Figure 6m-r, Table S15c |
| trait correlations | lmer(trait1 ~ trait2 + (1\|Site/Block)) | Table S3 |
| phenotypic on fruit (2019-2023), with family=binomial or truncated_poisson | glmmTMB(Fruit ~ Leaf lobing + Plant height + Flower width + Flowering time + Herkogamy + (1 \| Site/Block)) | Table S4 |
| herbivory between habitats | glmer(Herbivory ~ Habitat + (1\|Site/Block), family = binomial) | Lines 342-345 |
| herbivory and fitness | glmer(Seed ~ Herbivory + Habitat + Herbivory × Habitat + (1 \| Site/Block), family = poisson) | Lines 345-347 |
| Plasticity between habitats | glmmTMB(trait ~ Habitat + (1\|Site/Block)) | Tables S8, S9, S10 |
| plasticity between years | lmer(Phenotype ~ Habitat + Year + Habitat × Year + (1 \| Site/Block)) | Table S11 |
| snowpack and phenotypic traits, with family=binomial or truncated_poisson | glmmTMB(Fruit ~ Flowering time + Flower width + Plant height + Leaf lobing + Snowpack + Flowering time × Snowpack + Flower width × Snowpack + Plant height × Snowpack + Leaf lobing × Snowpack + (1 \| Site/Block) + (1 \| Year)) | Table S12 |
| differences in yearly selection gradients, with family=binomial or truncated_poisson | glmmTMB(Fruit ~ Flowering time + Year + Habitat + Flowering time × Year + Flowering time × Habitat + Year × Habitat + Flowering time × Year × Habitat + (1 \| Habitat:Site:Block)) | Tables S13, S14 |

**Table S7.**
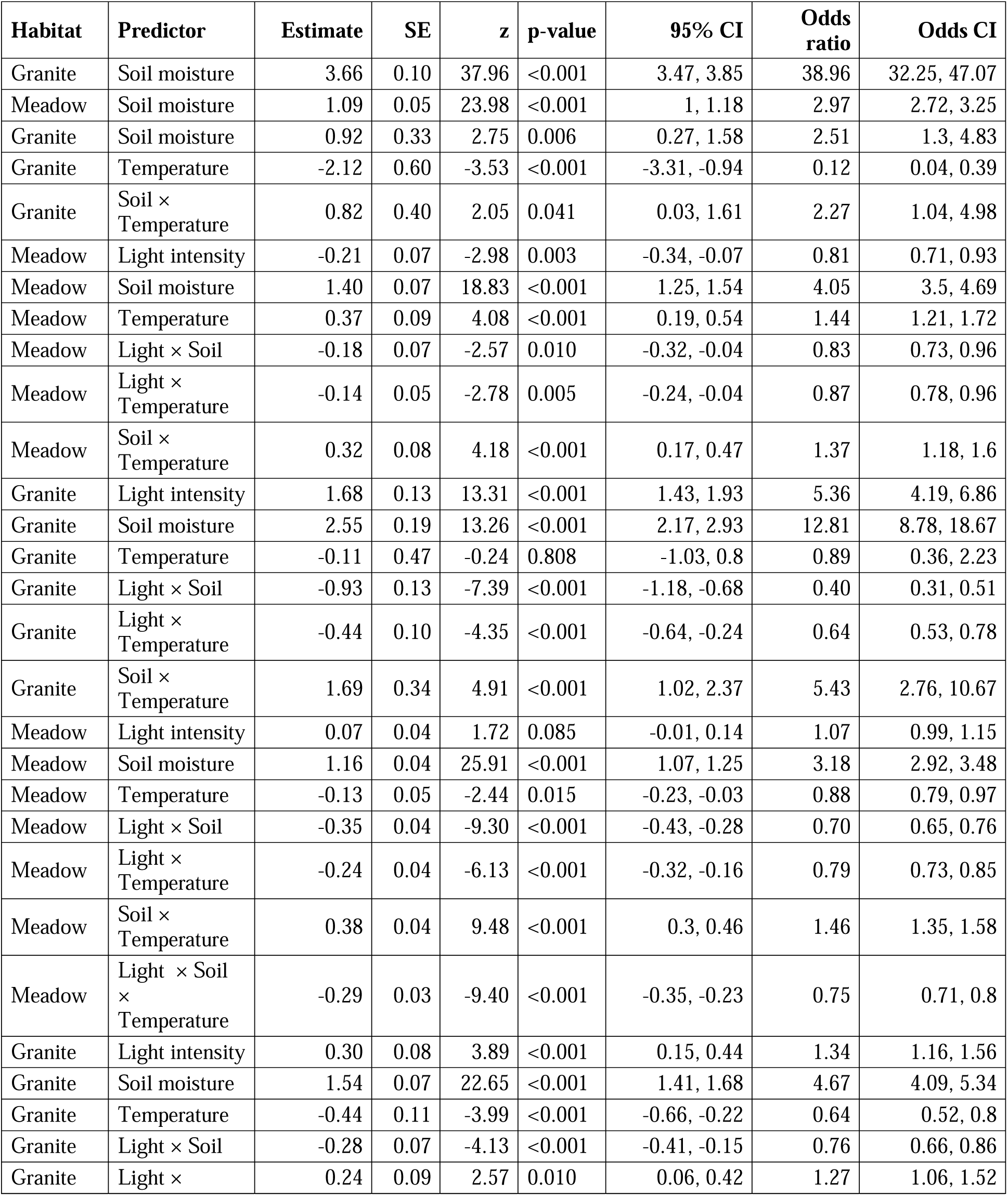
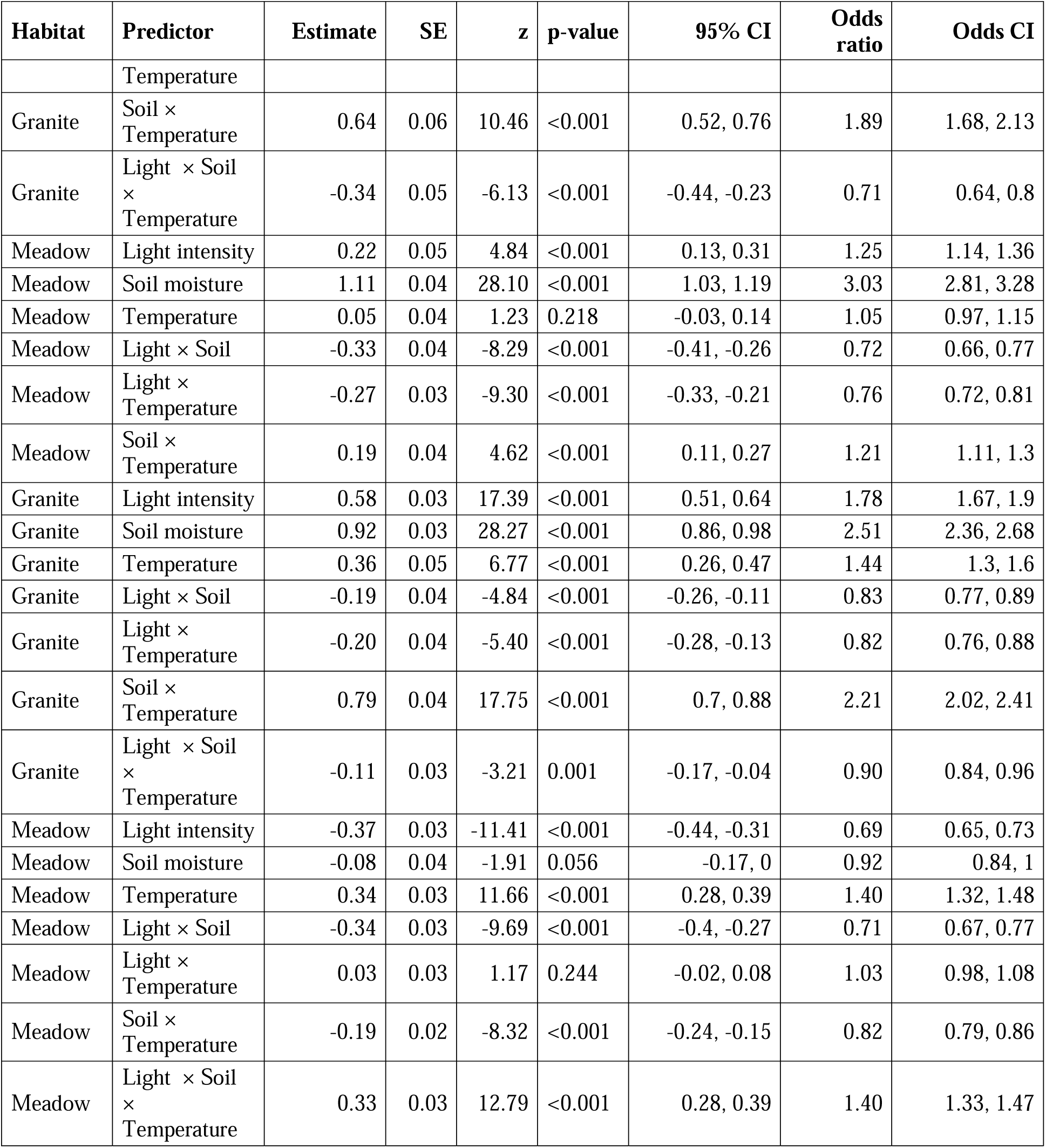
Best-fit models selected using AICc of the effect of environmental variables on overall plant survival in each habitat (Meadow or Granite) for 2013, 2019, 2021, 2022, 2023. Odds ratios represent the change in odds of interval survival associated with a one standard deviation increase in each predictor. Data from 2013 and 2019 is published in Ferris and Willis 2018 and Tataru et al. 2023 and re-analyzed here. significant p-values are in bold.

**Table S8.**
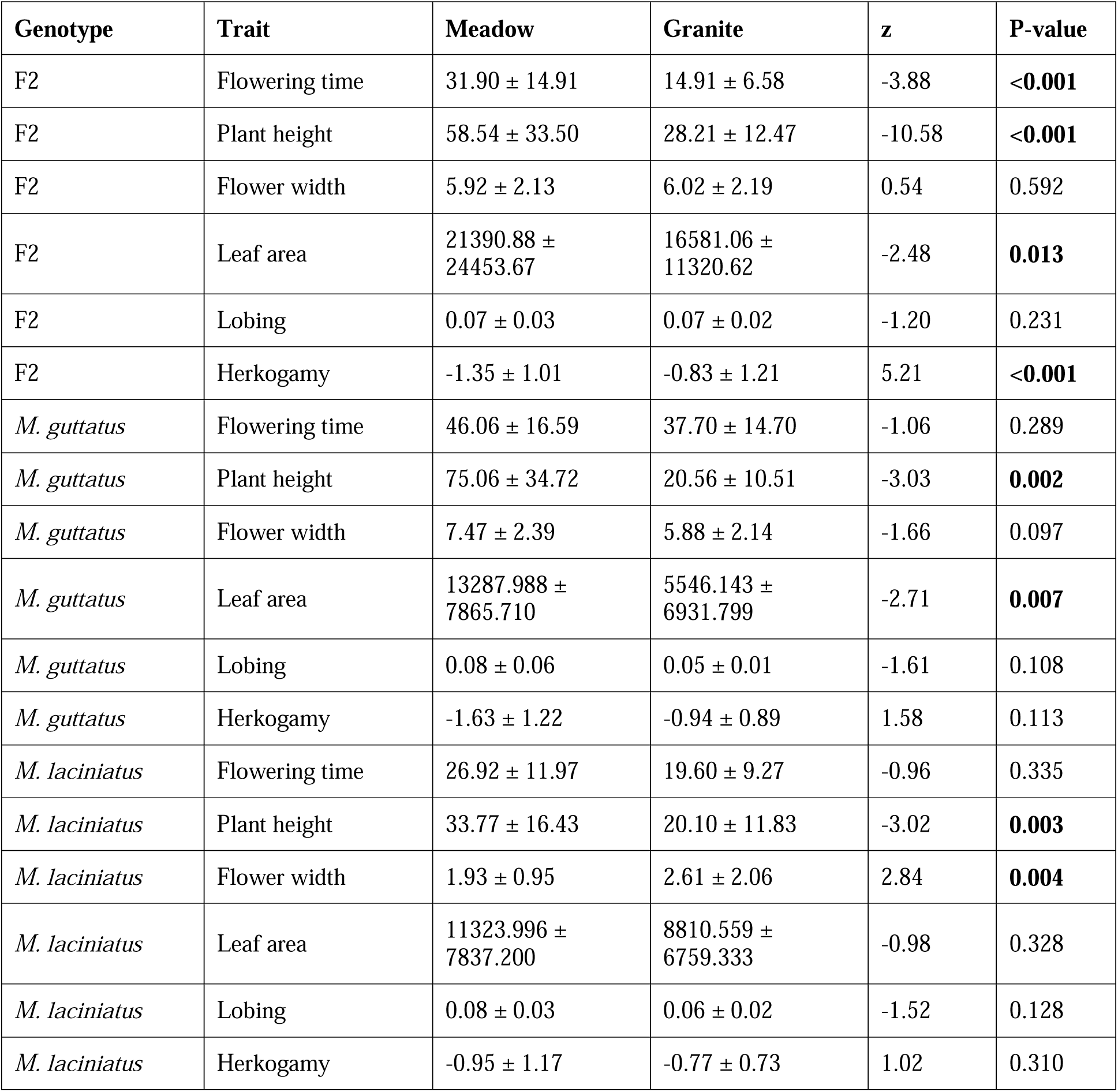
Trait differences between meadow and granite habitats for F2 hybrids, parental *M. guttatus*, and parental *M. laciniatus* from the 2021 experiment. Values are means ± SD. Habitat effects (e.g. plasticity) were evaluated using linear mixed models with Habitat as a fixed effect and Site and/or Block as random effects. Significant p-values are in bold. Trait means are reported in original units, although scaled values were used for model fitting of leaf area to improve model stability.

| Genotype | Trait | Meadow | Granite | z | P-value |
| --- | --- | --- | --- | --- | --- |
| F2 | Flowering time | 31.90 ± 14.91 | 14.91 ± 6.58 | -3.88 | **<0.001** |
| F2 | Plant height | 58.54 ± 33.50 | 28.21 ± 12.47 | -10.58 | **<0.001** |
| F2 | Flower width | 5.92 ± 2.13 | 6.02 ± 2.19 | 0.54 | 0.592 |
| F2 | Leaf area | 21390.88 ± 24453.67 | 16581.06 ± 11320.62 | -2.48 | **0.013** |
| F2 | Lobing | 0.07 ± 0.03 | 0.07 ± 0.02 | -1.20 | 0.231 |
| F2 | Herkogamy | -1.35 ± 1.01 | -0.83 ± 1.21 | 5.21 | **<0.001** |
| *M. guttatus* | Flowering time | 46.06 ± 16.59 | 37.70 ± 14.70 | -1.06 | 0.289 |
| *M. guttatus* | Plant height | 75.06 ± 34.72 | 20.56 ± 10.51 | -3.03 | **0.002** |
| *M. guttatus* | Flower width | 7.47 ± 2.39 | 5.88 ± 2.14 | -1.66 | 0.097 |
| *M. guttatus* | Leaf area | 13287.988 ± 7865.710 | 5546.143 ± 6931.799 | -2.71 | **0.007** |
| *M. guttatus* | Lobing | 0.08 ± 0.06 | 0.05 ± 0.01 | -1.61 | 0.108 |
| *M. guttatus* | Herkogamy | -1.63 ± 1.22 | -0.94 ± 0.89 | 1.58 | 0.113 |
| *M. laciniatus* | Flowering time | 26.92 ± 11.97 | 19.60 ± 9.27 | -0.96 | 0.335 |
| *M. laciniatus* | Plant height | 33.77 ± 16.43 | 20.10 ± 11.83 | -3.02 | **0.003** |
| *M. laciniatus* | Flower width | 1.93 ± 0.95 | 2.61 ± 2.06 | 2.84 | **0.004** |
| *M. laciniatus* | Leaf area | 11323.996 ± 7837.200 | 8810.559 ± 6759.333 | -0.98 | 0.328 |
| *M. laciniatus* | Lobing | 0.08 ± 0.03 | 0.06 ± 0.02 | -1.52 | 0.128 |
| *M. laciniatus* | Herkogamy | -0.95 ± 1.17 | -0.77 ± 0.73 | 1.02 | 0.310 |

**Table S9.** Trait differences between meadow and granite habitats for F3 hybrids, parental *M. guttatus*, and parental *M. laciniatus* from the 2022 experiment. Values are means ± SD. Habitat effects (e.g. plasticity) were evaluated using linear mixed models with Habitat as a fixed effect and Site and/or Block as random effects. Significant p-values are in bold. Trait means are reported in original units, although scaled values were used for model fitting of leaf area to improve model stability. No individuals of *M. guttatus* survived to flowering in granite habitat.

| Genotype | Trait | Meadow | Granite | z | P-value |
| --- | --- | --- | --- | --- | --- |
| F3 | Flowering time | 59.92 ± 12.75 | 44.19 ± 14.22 | -2.47 | **0.013** |
| F3 | Plant height | 54.89 ± 37.73 | 13.14 ± 6.97 | -2.66 | **0.008** |
| F3 | Flower width | 5.52 ± 2.26 | 3.34 ± 1.36 | -3.64 | **<0.001** |
| F3 | Leaf area | 1013.85 ± 1323.60 | 208.62 ± 226.80 | -1.92 | 0.054 |
| F3 | Lobing | 0.07 ± 0.02 | 0.09 ± 0.02 | 3.71 | **<0.001** |
| F3 | Herkogamy | -1.35 ± 0.91 | -0.92 ± 0.71 | 1.92 | 0.055 |
| *M. guttatus* | Flowering time | 72.66 ± 12.13 | — | — | — |
| *M. guttatus* | Plant height | 69.15 ± 33.42 | — | — | — |
| *M. guttatus* | Flower width | 7.73 ± 2.22 | — | — | — |
| *M. guttatus* | Leaf area | 1788.03 ± 1267.87 | 169.69 ± 79.60 | -2.23 | **0.026** |
| *M. guttatus* | Lobing | 0.05 ± 0.01 | 0.08 ± 0.02 | 2.57 | **0.010** |
| *M. guttatus* | Herkogamy | -2.40 ± 0.80 | — | — | — |
| *M. laciniatus* | Flowering time | 44.53 ± 11.01 | 34.65 ± 10.32 | -2.13 | **0.033** |
| *M. laciniatus* | Plant height | 15.61 ± 13.11 | 7.16 ± 4.06 | -2.65 | **0.008** |
| *M. laciniatus* | Flower width | 1.25 ± 0.92 | 1.22 ± 1.19 | -0.29 | 0.774 |
| *M. laciniatus* | Leaf area | 294.09 ± 285.43 | 97.56 ± 40.17 | -2.25 | 0.024 |
| *M. laciniatus* | Lobing | 0.08 ± 0.02 | 0.09 ± 0.03 | 1.81 | 0.070 |
| *M. laciniatus* | Herkogamy | -0.85 ± 0.55 | -1.01 ± 0.71 | -0.52 | 0.601 |

| Genotype | Trait | Meadow | Granite | z | P |
| --- | --- | --- | --- | --- | --- |
| F3 | Flowering time | 59.92 ± 12.75 | 44.19 ± 14.22 | -2.47 | **0.013** |
| F3 | Plant height | 54.89 ± 37.73 | 13.14 ± 6.97 | -2.66 | **0.008** |
| F3 | Flower width | 5.52 ± 2.26 | 3.34 ± 1.36 | -3.64 | **<0.001** |
| F3 | Leaf area | 1013.85 ± 1323.60 | 208.62 ± 226.80 | -1.92 | 0.054 |

| **Genotype** | **Trait** | **Meadow** | **Granite** | **z** | **P** |
| --- | --- | --- | --- | --- | --- |
| F3 | Lobing | 0.07 ± 0.02 | 0.09 ± 0.02 | 3.71 | **<0.001** |
| F3 | Herkogamy | -1.35 ± 0.91 | -0.92 ± 0.71 | 1.92 | 0.055 |
| *M. guttatus* | Flowering time | 72.66 ± 12.13 | NaN ± NA | — | — |
| *M. guttatus* | Plant height | 69.15 ± 33.42 | NaN ± NA | — | — |
| *M. guttatus* | Flower width | 7.73 ± 2.22 | NaN ± NA | — | — |
| *M. guttatus* | Leaf area | 1788.03 ± 1267.87 | 169.69 ± 79.60 | -2.23 | **0.026** |
| *M. guttatus* | Lobing | 0.05 ± 0.01 | 0.08 ± 0.02 | 2.57 | **0.010** |
| *M. guttatus* | Herkogamy | -2.40 ± 0.80 | NaN ± NA | — | — |
| *M. laciniatus* | Flowering time | 44.53 ± 11.01 | 34.65 ± 10.32 | -2.13 | **0.033** |
| *M. laciniatus* | Plant height | 15.61 ± 13.11 | 7.16 ± 4.06 | -2.65 | **0.008** |
| *M. laciniatus* | Flower width | 1.25 ± 0.92 | 1.22 ± 1.19 | -0.29 | 0.774 |
| *M. laciniatus* | Leaf area | 294.09 ± 285.43 | 97.56 ± 40.17 | -2.25 | 0.024 |
| *M. laciniatus* | Lobing | 0.08 ± 0.02 | 0.09 ± 0.03 | 1.81 | 0.070 |
| *M. laciniatus* | Herkogamy | -0.85 ± 0.55 | -1.01 ± 0.71 | -0.52 | 0.601 |

**Table S10.** Trait differences between meadow and granite habitats for RILs (F6 hybrids), parental *M. guttatus*, and parental *M. laciniatus* from the 2023 experiment. Values are means ± SD. Habitat effects (e.g. plasticity) were evaluated using linear mixed models with Habitat as a fixed effect and Site and/or Block as random effects. Significant p-values are in bold. Trait means are reported in original units, although scaled values were used for model fitting of leaf area to improve model stability.

| Genotype | Trait | Meadow | Granite | z | P-value |
| --- | --- | --- | --- | --- | --- |
| RIL | Flowering time | 43.04 ± 15.30 | 35.77 ± 19.67 | -4.63 | **<0.001** |
| RIL | Plant height | 91.71 ± 55.14 | 25.22 ± 14.81 | -2.15 | **0.032** |
| RIL | Flower width | 5.62 ± 2.23 | 4.68 ± 2.35 | -3.20 | **0.001** |
| RIL | Leaf area | 14342.24 ± 15722.42 | 6183.81 ± 9528.94 | -1.66 | 0.096 |
| RIL | Lobing | 0.08 ± 0.05 | 0.08 ± 0.06 | 0.48 | 0.633 |
| RIL | Herkogamy | -0.34 ± 1.40 | -0.40 ± 1.09 | -0.51 | 0.610 |
| *M. guttatus* | Flowering time | 52.94 ± 12.14 | 44.80 ± 15.38 | -3.74 | **<0.001** |
| *M. guttatus* | Plant height | 76.17 ± 34.94 | 35.82 ± 18.78 | -5.57 | **<0.001** |
| *M. guttatus* | Flower width | 9.92 ± 2.68 | 8.92 ± 3.42 | -1.38 | 0.169 |
| *M. guttatus* | Leaf area | 18876.10 ± 18833.71 | 12382.59 ± 14163.64 | -2.22 | **0.027** |
| *M. guttatus* | Lobing | 0.07 ± 0.03 | 0.08 ± 0.02 | 3.84 | **<0.001** |
| *M. guttatus* | Herkogamy | -0.66 ± 1.50 | -1.10 ± 1.40 | -1.62 | 0.106 |
| *M. laciniatus* | Flowering time | 24.28 ± 14.19 | 20.68 ± 18.36 | -1.02 | 0.307 |
| *M. laciniatus* | Plant height | 58.72 ± 34.45 | 21.16 ± 11.16 | -2.06 | 0.039 |
| *M. laciniatus* | Flower width | 3.01 ± 1.59 | 3.37 ± 1.44 | 2.15 | **0.032** |
| *M. laciniatus* | Leaf area | 17699.42 ± 16991.34 | 8788.10 ± 6537.97 | -1.25 | 0.210 |
| *M. laciniatus* | Lobing | 0.15 ± 0.08 | 0.11 ± 0.07 | -2.60 | **0.009** |
| *M. laciniatus* | Herkogamy | -0.36 ± 0.98 | 0.12 ± 0.92 | 2.72 | **0.007** |

**Table S11.**
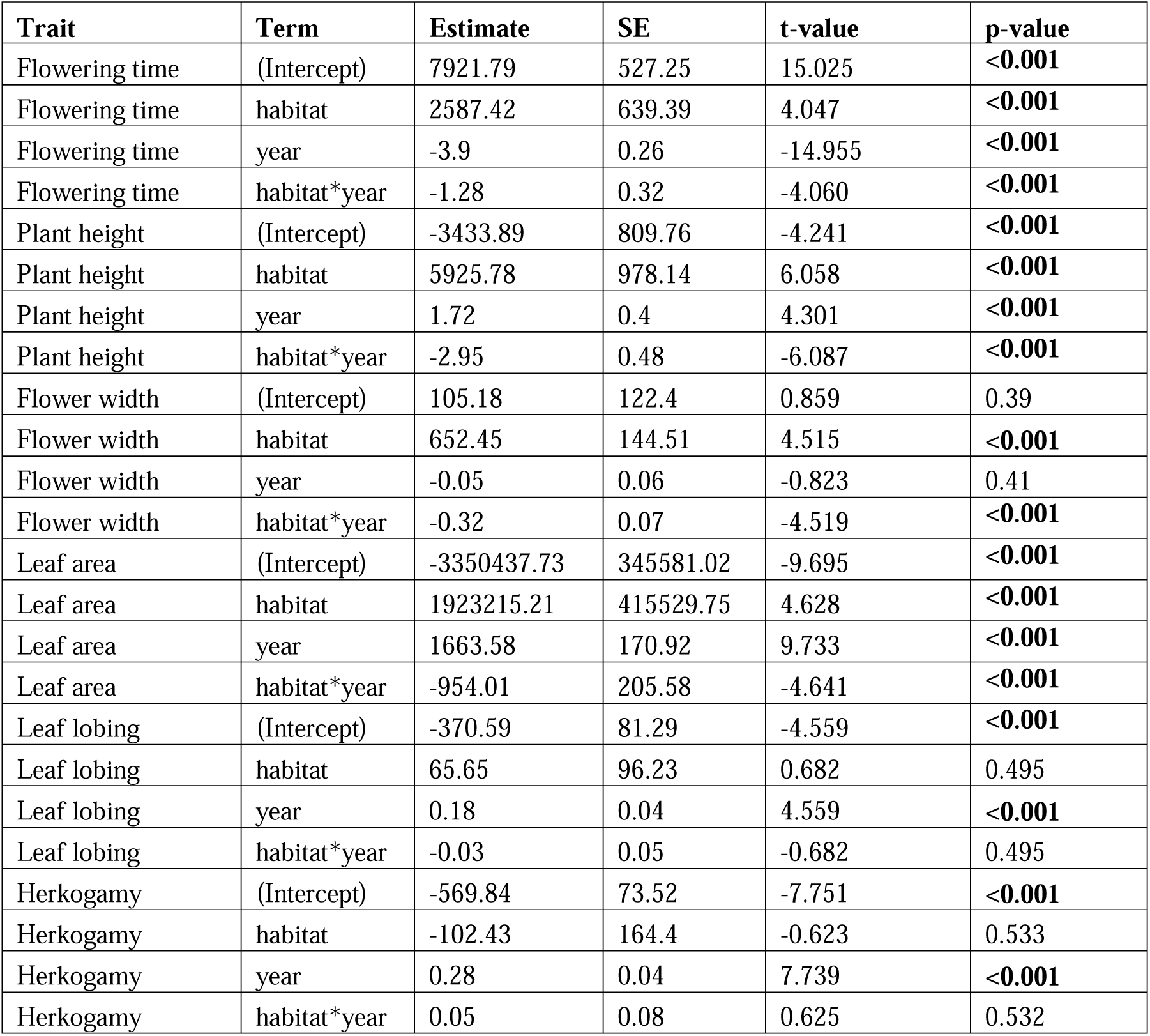
Generalized linear mixed models examining phenotypic plasticity between years and habitat. Models included the phenotypic trait as the dependent variable, habitat and year with their interaction as fixed effects and block nested within site as a random effect. Significant effects are indicated by *P < 0.05, **P < 0.01, ***P < 0.001. Significant p-values are in bold.

**Table S12.**
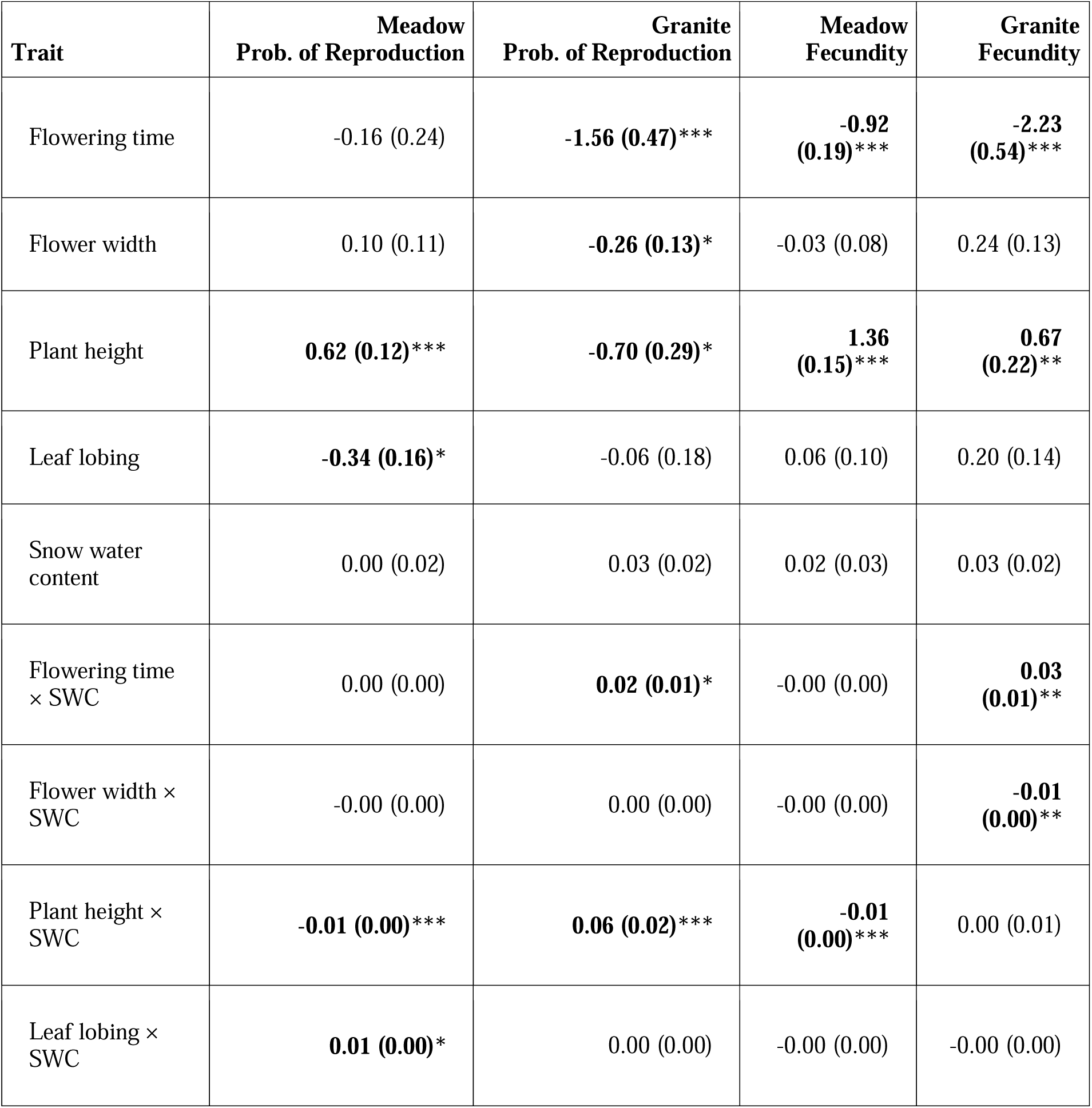
Effects of phenotypic traits, snow water content (SWC), and their interactions on the combined probability of survival and reproduction (binomial fitness) and fecundity (zero-truncated Poisson fitness) in meadow and granite habitats. Values are standardized selection gradients (β) with standard error (SE). Significant effects are indicated by *P < 0.05, **P < 0.01, ***P < 0.001 and in bold.

**Table S13.** Tests for temporal and spatial variation in phenotypic selection. Type III Wald χ² tests from generalized linear mixed models examining whether selection gradients differed among years and habitats (trait × year × habitat interaction). Significant interactions indicate that the relationship between a trait and fitness changed differently across habitats through time.

| Trait | $\chi^2$ | df | p-value |
| --- | --- | --- | --- |
| **Fecundity selection** |  |  |  |
| Flowering time | 21.14 | 4 | **<0.001** |
| Flower width | 9.09 | 4 | 0.059 |
| Plant height | 38.64 | 4 | **<0.001** |
| Leaf lobing | 3.80 | 4 | 0.434 |
| **Probability of reproduction** |  |  |  |
| Flowering time | 12.32 | 4 | **0.015** |
| Flower width | 12.10 | 4 | **0.017** |
| Plant height | 21.46 | 4 | **<0.001** |
| Leaf lobing | 3.12 | 4 | 0.539 |

**Table S14.** Estimated selection gradients (β) for each year × habitat combination, obtained as simple slopes from generalized linear mixed models including trait × year × habitat interactions. Selection gradients were estimated using emtrends, with separate conditional slopes reported for each habitat within each year. 95% confidence intervals are shown.

| Trait | Year | Habitat | Selection gradient | SE | 95% CI |
| --- | --- | --- | --- | --- | --- |
| **Fecundity** |  |  |  |  |  |
| Flowering time | 2013 | Meadow | -0.269 | 1.043 | -2.314, 1.776 |
| Flowering time | 2013 | Granite | -4.614 | 0.811 | -6.202, -3.025 |
| Flowering time | 2019 | Meadow | -1.277 | 0.201 | -1.671, -0.884 |
| Flowering time | 2019 | Granite | -1.231 | 0.295 | -1.809, -0.654 |
| Flowering time | 2021 | Meadow | -0.829 | 0.139 | -1.101, -0.557 |
| Flowering time | 2021 | Granite | -1.253 | 0.378 | -1.994, -0.512 |
| Flowering time | 2022 | Meadow | 0.018 | 0.251 | -0.474, 0.51 |
| Flowering time | 2022 | Granite | -0.954 | 2.363 | -5.585, 3.677 |
| Flowering time | 2023 | Meadow | -0.863 | 0.118 | -1.093, -0.632 |
| Flowering time | 2023 | Granite | -0.067 | 0.190 | -0.439, 0.306 |
| Plant height | 2013 | Meadow | 0.639 | 0.265 | 0.119, 1.159 |
| Plant height | 2013 | Granite | 0.745 | 0.141 | 0.469, 1.021 |
| Plant height | 2019 | Meadow | 1.860 | 0.292 | 1.288, 2.433 |
| Plant height | 2019 | Granite | 0.323 | 0.196 | -0.06, 0.706 |
| Plant height | 2021 | Meadow | 0.877 | 0.127 | 0.627, 1.127 |
| Plant height | 2021 | Granite | 2.373 | 0.695 | 1.011, 3.736 |
| Plant height | 2022 | Meadow | 1.253 | 0.216 | 0.829, 1.676 |
| Plant height | 2022 | Granite | 25.207 | 26.877 | -27.471, 77.886 |
| Plant height | 2023 | Meadow | 0.590 | 0.094 | 0.407, 0.774 |
| Plant height | 2023 | Granite | 3.771 | 0.822 | 2.16, 5.381 |
| **Probability of reproduction** |  |  |  |  |  |
| Flower width | 2013 | Meadow | -0.163 | 0.190 | -0.535, 0.209 |
| Flower width | 2013 | Granite | -0.005 | 0.134 | -0.269, 0.258 |
| Flower width | 2019 | Meadow | 0.238 | 0.154 | -0.063, 0.539 |
| Flower width | 2019 | Granite | -0.429 | 0.188 | -0.797, -0.061 |
| Flower width | 2021 | Meadow | 0.222 | 0.129 | -0.031, 0.476 |
| Flower width | 2021 | Granite | -0.341 | 0.132 | -0.599, -0.082 |
| Flower width | 2022 | Meadow | 0.084 | 0.160 | -0.229, 0.397 |
| Flower width | 2022 | Granite | -1.346 | 0.897 | -3.104, 0.413 |
| Flower width | 2023 | Meadow | -0.072 | 0.094 | -0.257, 0.113 |
| Flower width | 2023 | Granite | -0.183 | 0.164 | -0.506, 0.139 |
| Flowering time | 2013 | Meadow | 0.096 | 0.540 | -0.962, 1.155 |
| Flowering time | 2013 | Granite | -2.887 | 0.565 | -3.995, -1.78 |
| Flowering time | 2019 | Meadow | -0.560 | 0.213 | -0.978, -0.141 |
| Flowering time | 2019 | Granite | -1.092 | 0.919 | -2.893, 0.709 |
| Flowering time | 2021 | Meadow | -0.731 | 0.360 | -1.437, -0.024 |

| **Trait** | **Year** | **Habitat** | **Selection**<br>**gradient** | **SE** | **95% CI** |
| --- | --- | --- | --- | --- | --- |
| Flowering time | 2021 | Granite | -0.920 | 0.518 | -1.935, 0.095 |
| Flowering time | 2022 | Meadow | -0.069 | 0.253 | -0.564, 0.427 |
| Flowering time | 2022 | Granite | -0.050 | 0.851 | -1.717, 1.618 |
| Flowering time | 2023 | Meadow | 0.126 | 0.117 | -0.104, 0.356 |
| Flowering time | 2023 | Granite | -0.047 | 0.181 | -0.402, 0.308 |
| Plant height | 2013 | Meadow | 0.271 | 0.102 | 0.07, 0.471 |
| Plant height | 2013 | Granite | 0.022 | 0.159 | -0.291, 0.334 |
| Plant height | 2019 | Meadow | 0.255 | 0.569 | -0.86, 1.37 |
| Plant height | 2019 | Granite | 0.097 | 0.779 | -1.429, 1.624 |
| Plant height | 2021 | Meadow | 1.285 | 0.528 | 0.25, 2.319 |
| Plant height | 2021 | Granite | 0.467 | 1.121 | -1.729, 2.664 |
| Plant height | 2022 | Meadow | 0.692 | 0.325 | 0.054, 1.329 |
| Plant height | 2022 | Granite | -7.152 | 7.486 | -21.825, 7.52 |
| Plant height | 2023 | Meadow | -0.474 | 0.139 | -0.747, -0.202 |
| Plant height | 2023 | Granite | 3.707 | 0.962 | 1.821, 5.592 |

**Table S15.**
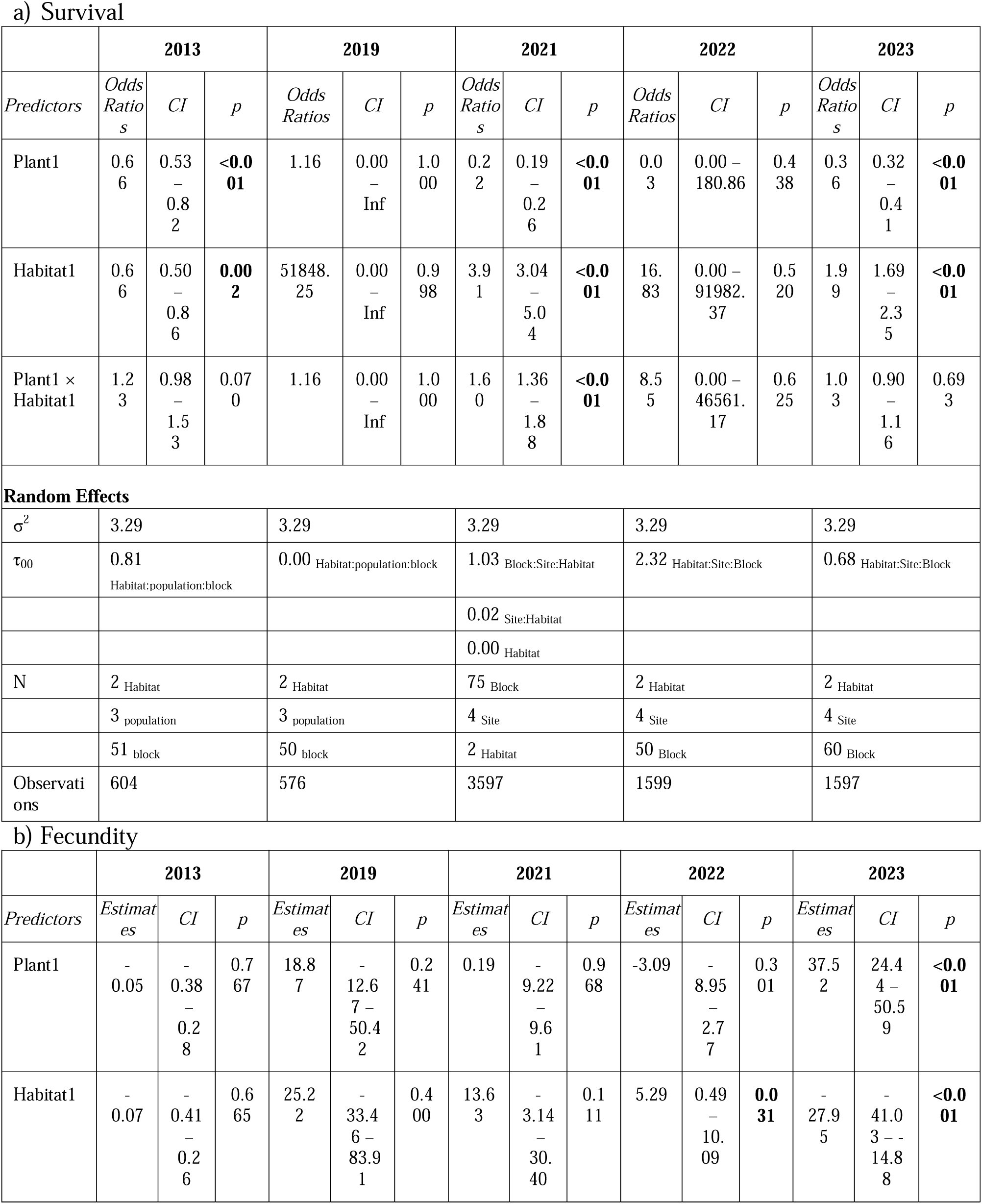
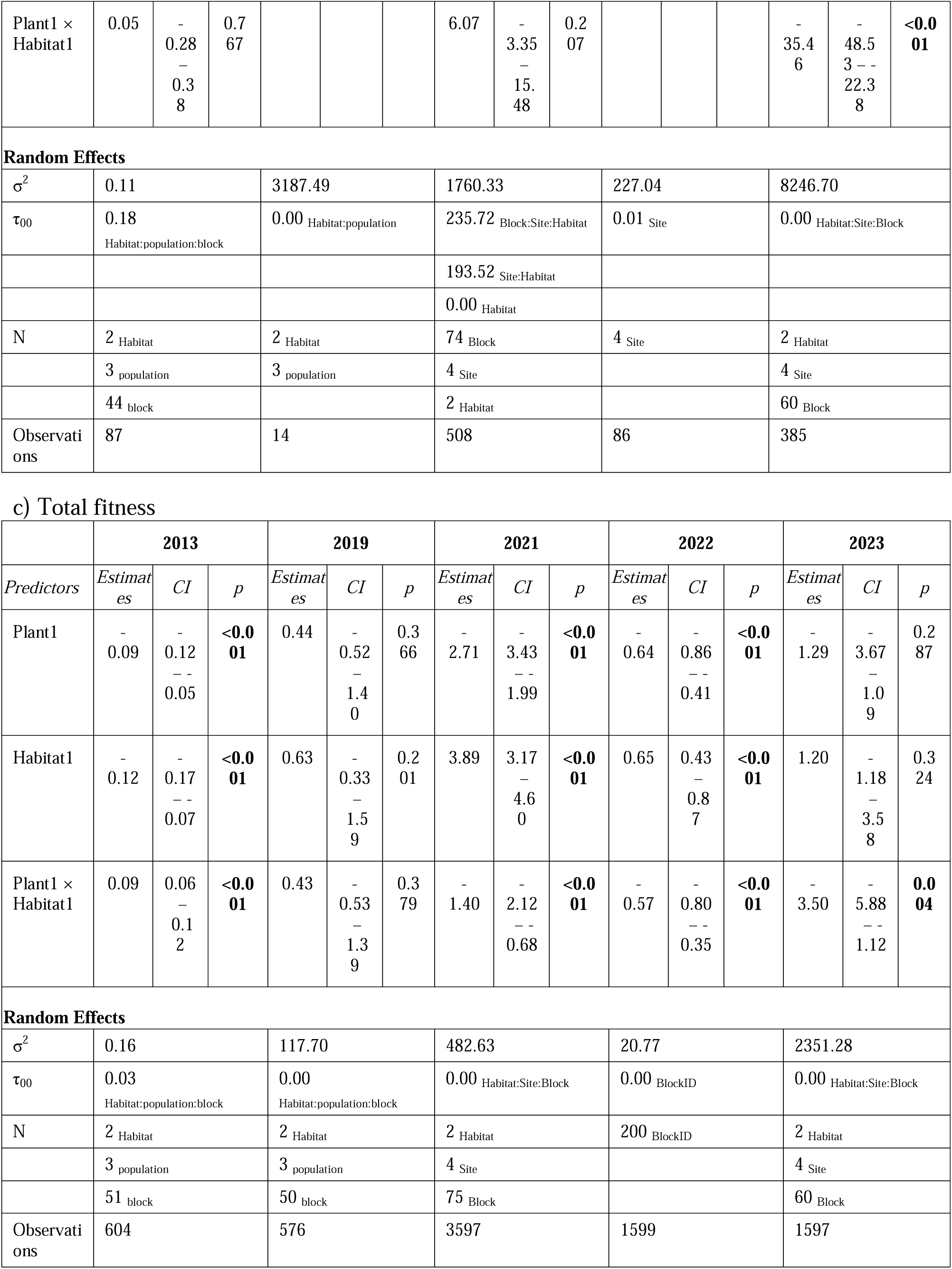
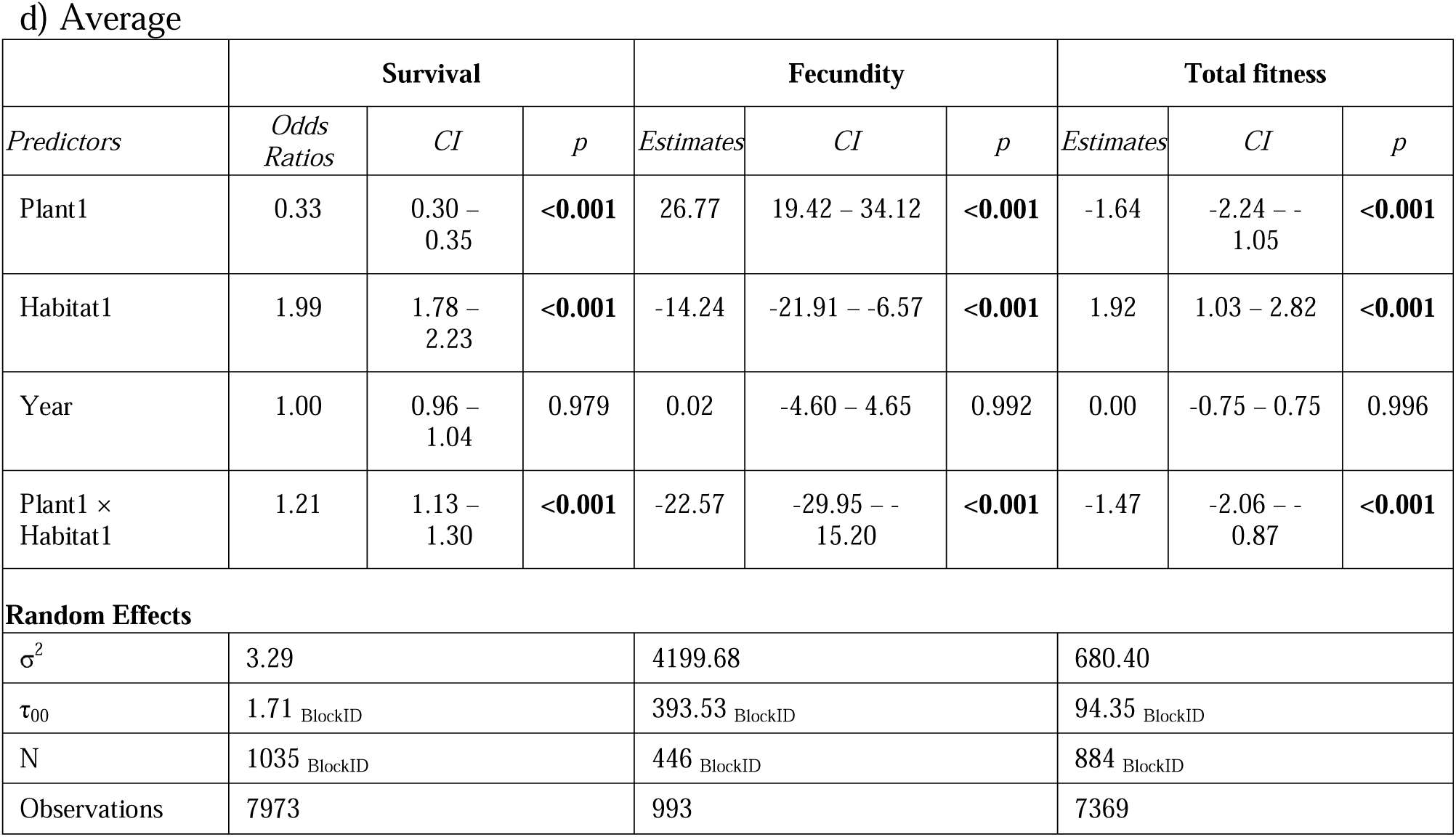
Significance of genotype (G; *Mimulus guttaus* or *M. laciniatus* species), environment (E; meadow or granite habitat), and genotype by environment (G×E) interactions for each fitness term (a-c) in each year and d) of all years averaged. Significant p-values are in bold (p < 0.05).

## REFERENCES

1. Aarseen, L. W., and D. R. Taylor. 1992. Fecundity allocation in herbaceous plants. Oikos 65:225–232.

2. Aarssen, L. W., and C. Y. Jordan. 2001. Between-species patterns of covariation in plant size, seed size, and fecundity in monocarpic herbs. Ecoscience 8:471–477.

3. Ågren, J., C. G. Oakley, S. Lundemo, and D. W. Schemske. 2017. Adaptive divergence in flowering time among natural populations of *Arabidopsis thaliana* : Estimates of selection and QTL mapping. Evolution 71:550–564.

4. Anderson, J. T. 2016. Plant fitness in a rapidly changing world. New Phytologist 210:81–87.

5. Anderson, J. T., V. M. Eckhart, and M. A. Geber. 2015a. Experimental studies of adaptation in Clarkia xantiana: III: Phenotypic selection across a subspecies border. Evolution 69:2249–2261.

6. Anderson, J. T. 2015b. Experimental studies of adaptation in Clarkia xantiana: III: Phenotypic selection across a subspecies border. Evolution 69:2249–2261.

7. Anderson, J. T., and S. M. Wadgymar. 2020. Climate change disrupts local adaptation and favours upslope migration. Ecology Letters 23:181–192.

8. Anderson, J. T., J. H. Willis, and T. Mitchell-Olds. 2011. Evolutionary genetics of plant adaptation. Trends in Genetics 27:258–266.

9. Awadalla, P., and K. Ritland. 1997. Microsatellite variation and evolution in the Mimulus guttatus species complex with contrasting mating systems. Molecular Biology and Evolution 14:1023–1034.

10. Bailey, L. D., and M. van de Pol. 2016. Tackling extremes: Challenges for ecological and evolutionary research on extreme climatic events. Journal of Animal Ecology 85:85–96.

11. Bartoń, K. 2018. MuMIn: Multi-Model Inference. R package version 1.42.1. https://CRAN.R-project.org/package=MuMIn.

12. Bates, D., M. Mächler, B. M. Bolker, and S. C. Walker. 2015. Fitting linear mixed-effects models using lme4. Journal of Statistical Software 67:1–48.

13. Bennett, A. F., and R. E. Lenski. 2007. An experimental test of evolutionary trade-offs during temperature adaptation. Proceedings of the National Academy of Sciences of the United States of America 104:8649–8654.

14. Bradshaw, H. D., S. M. Wilbert, K. G. Otto, and D. W. Schemske. 1995. Genetic mapping of floral traits associated with reproductive isolation in monkeyflowers (Mimulus). Nature 376:762–765.

15. Breheny, P., and W. Burchett. 2017. Visualization of Regression Models Using visreg. The R Journal 9:56–71.

16. Brooks, M. E., K. Kristensen, K. J. van Benthem, A. Magnusson, C. W. Berg, A. Nielsen, H. J. Skaug, et al. 2017. Modeling Zero-Inflated Count Data With glmmTMB. bioRxiv preprint. bioRxiv.

17. Brunet, J. 2009. Pollinators of the Rocky Mountain columbine: Temporal variation, functional groups and associations with floral traits. Annals of Botany 103:1567–1578.

18. Campbell, D. R., and J. M. Powers. 2015. Natural selection on floral morphology can be influenced by climate. Proceedings of the Royal Society B: Biological Sciences 282.

19. Campbell-Staton, S. C., Z. A. Cheviron, N. Rochette, J. Catchen, J. B. Losos, and S. V. Edwards. 2017. Winter storms drive rapid phenotypic, regulatory, and genomic shifts in the green anole lizard. Science 357:495–498.

20. Chapurlat, E., I. Le Roncé, J. Ågren, and N. Sletvold. 2020. Divergent selection on flowering phenology but not on floral morphology between two closely related orchids. Ecology and Evolution 10:5737–5747.

21. Coltman, D. W., J. A. Smith, D. R. Bancroft, J. Pilkington, A. D. C. MacColl, T. H. Clutton-Brock, and J. M. Pemberton. 1999. Density-dependent variation in lifetime breeding success and natural and sexual selection in Soay rams. American Naturalist 154:730–746.

22. Coyne, J. A., and H. A. Orr. 2004. Speciation. Sinauer Associates. Sinauer Associates, Sunderland, Massachusetts.

23. Crespi, B. J. 2000. The evolution of maladaptation. Heredity 84:623–629.

24. DeMarche, M. L., K. J. Rice, and J. P. Sexton. 2013. Niche partitioning between close relatives suggests trade-offs between adaptation to local environments and competition. Ecology and Evolution 3:512–522.

25. Dittmar, E. L., and D. W. Schemske. 2023. Temporal Variation in Selection Influences Microgeographic Local Adaptation. American Naturalist 202:471–485.

26. Ferris, K. G. 2019. Endless forms most functional: uncovering the role of natural selection in the evolution of leaf shape. American Journal of Botany 106:1532–1535.

27. Ferris, K. G., L. L. Barnett, B. K. Blackman, and J. H. Willis. 2017a. The genetic architecture of local adaptation and reproductive isolation in sympatry within the Mimulus guttatus species complex. Molecular Ecology 26:208–224.

28. Ferris, K. G., L. L. Barnett, B. K. Blackman, and J. H. Willis. 2017b. The genetic architecture of local adaptation and reproductive isolation in sympatry within the Mimulus guttatus species complex. Molecular Ecology 26:208–224.

29. Ferris, K. G., T. Rushton, A. B. Greenlee, K. Toll, B. K. Blackman, and J. H. Willis. 2015. Leaf shape evolution has a similar genetic architecture in three edaphic specialists within the Mimulus guttatus species complex. Annals of Botany 116:213–223.

30. Ferris, K. G., J. P. Sexton, and J. H. Willis. 2014. Speciation on a local geographic scale: The evolution of a rare rock outcrop specialist in Mimulus. Philosophical Transactions of the Royal Society B: Biological Sciences 369:27–29.

31. Ferris, K. G., and J. H. Willis. 2018. Differential adaptation to a harsh granite outcrop habitat between sympatric Mimulus species. Evolution 72:1225–1241.

32. Fishman, L., A. L. Sweigart, A. M. Kenney, and S. Campbell. 2014. Major quantitative trait loci control divergence in critical photoperiod for flowering between selfing and outcrossing species of monkeyflower (Mimulus). New Phytologist 201:1498–1507.

33. Friedman, J., and J. H. Willis. 2013. Major QTLs for critical photoperiod and vernalization underlie extensive variation in flowering in the Mimulus guttatus species complex. New Phytologist 199:571–583.

34. Fry, J. D. 1996. The Evolution of Host Specialization: Are Trade-Offs Overrated? The American Naturalist 148:S84–S107.

35. Galen, C. 2000. High and dry: Drought stress, sex-allocation trade-offs, and selection on flower size in the alpine wildflower Polemonium viscosum (Polemoniaceae). American Naturalist 156:72–83.

36. Ghalambor, C. K., K. L. Hoke, E. W. Ruell, E. K. Fischer, D. N. Reznick, and K. A. Hughes. 2015. Non-adaptive plasticity potentiates rapid adaptive evolution of gene expression in nature. Nature 525:372–375.

37. Gosden, T. P., J. T. Waller, and E. I. Svensson. 2015. Asymmetric isolating barriers between different microclimatic environments caused by low immigrant survival. Proceedings of the Royal Society B: Biological Sciences 282.

38. Grant, B. R., and P. R. Grant. 1993. Evolution of Darwin’s Finches Caused by a Rare Climatic Event. Proceedings of the Royal Society B: Biological Sciences 251:111–117.

39. Groh, J. S., and G. Coop. 2024. The temporal and genomic scale of selection following hybridization. Proceedings of the National Academy of Sciences of the United States of America 121:1–11.

40. Hall, M. C., and J. H. Willis. 2006. Divergent Selection on Flowering Time Contributes To Local Adaptation in Mimulus Guttatus Populations. Evolution 60:2466.

41. Harder, L. D., and S. D. Johnson. 2009. Darwin’s beautiful contrivances: evolutionary and functional evidence for floral adaptation. New Phytologist 183:530–545.

42. Hastings, A. 1983. Can spatial variation alone lead to selection for dispersal? Theoretical Population Biology 24:244–251.

43. Hereford, J. 2009. A quantitative survey of local adaptation and fitness trade-offs. American Naturalist 173:579–588.

44. Johnson, O. L., R. Tobler, J. M. Schmidt, and C. D. Huber. 2023. Fluctuating selection and the determinants of genetic variation. Trends in Genetics 39:491–504.

45. Kawecki, T. J., and D. Ebert. 2004. Conceptual issues in local adaptation. Ecology Letters 7:1225–1241.

46. Keep, T., S. Rouet, J. L. Blanco-Pastor, P. Barre, T. Ruttink, K. J. Dehmer, M. Hegarty, et al. 2021. Inter-annual and spatial climatic variability have led to a balance between local fluctuating selection and wide-range directional selection in a perennial grass species. Annals of Botany 128:357–369.

47. Kelly, J. K. 2022. The genomic scale of fluctuating selection in a natural plant population. Evolution Letters 6:506–521.

48. Kenney, A. M., and A. L. Sweigart. 2016. Reproductive isolation and introgression between sympatric Mimulus species. Molecular ecology 25:2499–2517.

49. Kingsolver, J. G., and S. E. Diamond. 2011. Phenotypic selection in natural populations: What limits directional selection? American Naturalist 177:346–357.

50. Kingsolver, J. G., and D. W. Pfennig. 2004. Individual-level selection as a cause of cope’s rule of phyletic size increase. Evolution 58:1608–1612.

51. Kooyers, N. J., J. M. Colicchio, A. B. Greenlee, E. Patterson, N. T. Handloser, and B. K. Blackman. 2019. Lagging adaptation to climate supersedes local adaptation to herbivory in an annual monkeyflower. American Naturalist 194:541–557.

52. Kooyers, N. J., A. B. Greenlee, J. M. Colicchio, M. Oh, and B. K. Blackman. 2015. Replicate altitudinal clines reveal that evolutionary flexibility underlies adaptation to drought stress in annual Mimulus guttatus. New Phytologist 206:152–165.

53. Krizek, B. A., and J. T. Anderson. 2013. Control of flower size. Journal of Experimental Botany 64:1427–1437.

54. Lande, R., and S. J. Arnold. 1983. The Measurement of Selection on Correlated Characters. Evolution 37:1210–1226.

55. Lane, J. E., L. E. B. Kruuk, A. Charmantier, J. O. Murie, and F. S. Dobson. 2012. Delayed phenology and reduced fitness associated with climate change in a wild hibernator. Nature 489:554–557.

56. Latreille, A. C., and C. Pichot. 2017. Local-scale diversity and adaptation along elevational gradients assessed by reciprocal transplant experiments: lack of local adaptation in silver fir populations. Annals of Forest Science 74.

57. Lee, G., B. J. Sanderson, T. J. Ellis, B. P. Dilkes, J. K. McKay, J. Ågren, and C. G. Oakley. 2024. A large-effect fitness trade-off across environments is explained by a single mutation affecting cold acclimation. Proceedings of the National Academy of Sciences 121.

58. Leidinger, L., D. Vedder, and J. S. Cabral. 2021. Temporal environmental variation may impose differential selection on both genomic and ecological traits. Oikos 130:1100–1115.

59. Leimu, R., and M. Fischer. 2008. A meta-analysis of local adaptation in plants. PLoS ONE 3:1–8.

60. Levins, R. 1968. Evolution in Changing Environments. Princeton University Press, Princeton, New Jersey.

61. Love, J. M., and K. G. Ferris. 2024. Local adaptation to an altitudinal gradient: The interplay between mean phenotypic trait variation and phenotypic plasticity in Mimulus laciniatus. Perspectives in Plant Ecology, Evolution and Systematics 63:125795.

62. Lowry, D. B., M. C. Hall, D. E. Salt, and J. H. Willis. 2009. Genetic and physiological basis of adaptive salt tolerance divergence between coastal and inland Mimulus guttatus. New Phytologist 183:776–788.

63. Lowry, D. B., R. C. Rockwood, and J. H. Willis. 2008. Ecological reproductive isolation of coast and inland races of Mimulus guttatus. Evolution 62:2196–2214.

64. Magnusson, A., H. J. Skaug, A. Nielsen, C. W. Berg, K. Kristensen, M. Maechler, K. J. van Bentham, et al. 2017. glmmTMB: Generalized Linear Mixed Models using Template Model Builder. R package version 0.1.3.

65. Mantel, S. J., and A. L. Sweigart. 2019. Divergence in drought-response traits between sympatric species of Mimulus. Ecology and Evolution 9:10291–10304.

66. Mills, L. S., M. Zimova, J. Oyler, S. Running, J. T. Abatzoglou, and P. M. Lukacs. 2013. Camouflage mismatch in seasonal coat color due to decreased snow duration. Proceedings of the National Academy of Sciences of the United States of America 110:7360–7365.

67. Mojica, J. P., and J. K. Kelly. 2010. Viability selection prior to trait expression is an essential component of natural selection. Proceedings of the Royal Society B: Biological Sciences 277:2945–2950.

68. Mojica, J. P., Y. W. Lee, J. H. Willis, and J. K. Kelly. 2012. Spatially and temporally varying selection on intrapopulation quantitative trait loci for a life history trade-off in Mimulus guttatus. Molecular Ecology 21:3718–3728.

69. Monnahan, P. J., and J. K. Kelly. 2015. Naturally segregating loci exhibit epistasis for fitness. Biology Letters 11.

70. Nagy, E. S. 1997. Selection for native characters in hybrids between two locally adapted plant subspecies. Evolution 51:1469–1480.

71. Nicotra, A. B., A. Leigh, C. K. Boyce, C. S. Jones, K. J. Niklas, D. L. Royer, and H. Tsukaya. 2011a. The evolution and functional significance of leaf shape in the angiosperms. Functional Plant Biology 38:535–552.

72. Nicotra, A. B., A. Leigh, C. K. Boyce, C. S. Jones, K. J. Niklas, D. L. Royer, and H. Tsukaya. 2011b. The evolution and functional significance of leaf shape in the angiosperms. Functional Plant Biology 38:535–552.

73. Oakley, C. G., D. W. Schemske, J. K. McKay, and J. Ågren. 2023. Ecological genetics of local adaptation in Arabidopsis: An 8-year field experiment. Molecular Ecology 32:4570–4583.

74. Pemberton, J. M., L. E. B. Kruuk, and T. Clutton-Brock. 2022. The Unusual Value of Long-Term Studies of Individuals: The Example of the Isle of Rum Red Deer Project. Annual Review of Ecology, Evolution, and Systematics 53:327–351.

75. Pinheiro, J., and D. Bates. 2000. Mixed-Effects Models in S and S-PLUS. Springer, New York.

76. Pinheiro, J., D. Bates, and R Core Team. 2023. nlme: Linear and Nonlinear Mixed Effects Models. R package version 3.1–163.

77. Ramsey, J., H. D. Bradshaw, and D. W. Schemske. 2003. Components of reproductive isolation between the monkeyflowers Mimulus lewisii and M. cardinalis (Phrymaceae). Evolution 57:1520–1534.

78. Rundle, H. D., and P. Nosil. 2005. Ecological speciation. Ecology Letters 8:336–352.

79. Schiestl, F. P., and P. M. Schluter. 2009. Floral isolation, specialized pollination, and pollinator behavior in orchids. Annual Review of Entomology 54:425–446.

80. Schluter, D. 2009. Evidence for ecological speciation and its alternative. Science 323:737–741.

81. Schneider, C. A., W. S. Rasband, and K. W. Eliceiri. 2012. NIH Image to ImageJ: 25 years of Image Analysis. Nature Methods 9:671–675.

82. Siepielski, A. M., J. D. Dibattista, and S. M. Carlson. 2009. It’s about time: The temporal dynamics of phenotypic selection in the wild. Ecology Letters 12:1261–1276.

83. Stearns, S. 1992. The evolution of life histories. Oxford University Press, Oxford, UK.

84. Tataru, D., M. De Leon, S. Dutton, F. Machado-Perez, A. Rendahl, and K. G. Ferris. 2024. Fluctuating selection in a Monkeyflower hybrid zone. bioRxiv 1–33.

85. Tataru, D., E. C. Wheeler, and K. G. Ferris. 2023. Spatially and temporally varying selection influence species boundaries in two sympatric Mimulus. Proceedings of the Royal Society B: Biological Sciences 290.

86. Toll, K., and J. H. Willis. 2018. Hybrid inviability and differential submergence tolerance drive habitat segregation between two congeneric monkeyflowers. Ecology 99:2776–2786.

87. Troth, A., J. R. Puzey, R. S. Kim, J. H. Willis, and J. K. Kelly. 2018. Selective trade-offs maintain alleles underpinning complex trait variation in plants. Science 361:475–478.

88. Venail, J., A. Dell’Olivo, and C. Kuhlemeier. 2010. Speciation genes in the genus Petunia. Philosophical Transactions of the Royal Society B: Biological Sciences 365:461–468.

89. Via, S., and R. Lande. 1985. Genotype-environment interaction and the evolution of phenotypic plasticity. Evolution 39:505–522.

90. Vickery, R. K. 1964. Barriers to gene exchange between members of the Mimulus guttatus complex (Scrophulariaceae). Evolution 18:52.

91. Wadgymar, S. M., S. C. Daws, and J. T. Anderson. 2017. Integrating viability and fecundity selection to illuminate the adaptive nature of genetic clines. Evolution Letters 1:26–39.

92. Wilczek, A. M., M. D. Cooper, T. M. Korves, and J. Schmitt. 2014. Lagging adaptation to warming climate in Arabidopsis thaliana. Proceedings of the National Academy of Sciences of the United States of America 111:7906–7913.

93. Willis, J. 1999. The role of genes of large effect on inbreeding depression in Mimulus guttatus. Evolution 53:1678–1691.

94. Wright, S. I., S. Kalisz, and T. Slotte. 2013. Evolutionary consequences of self-fertilization in plants. Proceedings of the Royal Society B: Biological Sciences 280.

95. Wu, C. A., D. B. Lowry, A. M. Cooley, K. M. Wright, Y. W. Lee, and J. H. Willis. 2008. Mimulus is an emerging model system for the integration of ecological and genomic studies. Heredity 100:220–230.

